# Localized translation of *erm-1* contributes to ERM-1 function in the *C. elegans* embryo

**DOI:** 10.1101/2024.09.23.614538

**Authors:** Elise van der Salm, Esther Koelewijn, Erica van der Maas, Max Eeken, Suzan Ruijtenberg

## Abstract

Translation of mRNAs into proteins is a key step in decoding the information stored in the genome. Localized translation ensures that proteins are expressed where needed, which is important for cell-specific protein expression, the establishment of cellular protein gradients, and the creation of protein hotspots within different cellular compartments. Although localized translation is believed to be important for cell fate determination and organismal development, our understanding of localized translation in the context of living animals is limited, as few methods that allow direct visualization and measurement of translation exist. We adapted the SunTag-based single-molecule translation imaging system for use in Caenorhabditis elegans and showed the dynamics and importance of localized *erm-1* translation during development. We found *erm-1* translation to be enriched at the plasma membrane, overlapping with the localization and function of the encoded membrane-cytoskeleton linker ERM-1. Re-localizing *erm-1* translation to nuclear pores disrupts the function of ERM-1 protein, particularly its role in linking the actin cytoskeleton to the membrane. Our work demonstrates the power of translation imaging and highlights the importance of localized translation in *C. elegans* development.

## Introduction

Expressing the correct complement of proteins in the right cells at the correct time is crucial for the development of multicellular organisms. Even small changes in expression of a single protein can lead to malfunctioning tissues and disease (Buszczak et al., 2014; Fero et al., 1998; Overton et al., 2014). A key step in achieving accurate protein levels is regulation at the level of translation, the process of decoding the information from mRNAs into amino-acid sequences to synthesize functional proteins. The regulation of translation allows for temporal and spatially controlled gene expression regulation after transcripts are produced. This is particularly important during developmental transitions and cellular differentiation, when rapid changes in the proteome are required (Sonneveld et al., 2020). As such, several developmental pathways have been identified, that control ‘where’, ‘when’ and ‘how efficiently’ mRNAs are translated into proteins.

Localized translation, defined as the synthesis of proteins in specific cells or at specific subcellular regions, is one way to ensure that proteins are expressed exactly where they are needed (Das et al., 2021). It contributes to the establishment of gradients across tissues, the generation of cell type-specific protein expression profiles, and the differentiation of protein composition among different cellular compartments. Localized translation can either be regulated at a global level, influencing all mRNAs within the cell, or at an mRNA-specific level. At the global level, localizing crucial components of the translation machinery can enhance translation efficiency at specific places in the cell. For example, ribosome recruitment to the apical side of intestinal cells in mice results in apically enhanced translation efficiencies upon refeeding, thereby boosting nutrient absorption (Silva et al., 2022). mRNA-specific localized translation can be achieved by inhibiting or promoting translation of specific mRNA molecules at particular sites. For example, during early development of *Drosophila melanogaster, caudal* mRNA is translationally active only at the posterior side of the embryo due to the asymmetric distribution of the translational repressor Bicoid (Niessing et al., 2002). Similarly, in the early *Caenorhabditis elegans* embryo, *glp-1* mRNAs are present throughout the embryo, whereas the encoded developmental regulator GLP-1 Notch proteins are present only in the anterior precursor cells. This regulation of GLP-1 expression is mediated by cell-specific combinations of RNA binding proteins that promote *glp-1* translation in the anterior but prevent *glp-1* translation in the posterior cells (Evans et al., 1994; Farley & Ryder, 2012; Ogura et al., 2003). Localized translation has also been extensively observed in neurons, where many mRNAs that are needed for the establishment and maintenance of synaptic connections are specifically translated in distal dendrites and axons (Dalla Costa et al., 2021; Holt et al., 2019). Moreover, *actin* is particularly translated at the leading edge of migrating fibroblasts (Huttelmaier et al., 2005), and multiple centrosome-associated proteins are found to be translated at the centrosomes in different cell types (Safieddine et al., 2021; Sepulveda et al., 2018). As such, localized translation corresponds to protein levels and localization, and prevents the accumulation of proteins in undesired places of the cell. In addition, localized translation in the right micro-environment has been suggested to be important for the maturation of newly synthesized proteins, facilitating posttranslational modifications and fast assembly of protein complexes (Das et al., 2021).

Localized translation is often intertwined with mRNA localization; if mRNAs are localized to specific sites in the cell, it is highly likely that translation will follow that pattern. However, the opposite has also been observed, where mRNA localization is dependent on and guided by translation (Das et al., 2021). In the latter case, the localization signal is not provided by the mRNA sequence, but a sequence in the nascent (poly)peptide. This is best described for secreted proteins, which have a nascent signal peptide that is immediately recognized by a signal recognition particle, leading to the inhibition of translation elongation, and translocation to the endoplasmic reticulum where translation elongation resumes (Reid & Nicchitta, 2015). More recently, translation-dependent mRNA localization was shown to occur for many mRNAs that are translated in the cytosol. Here, the mRNA-ribosome complex is probably localized by larger nascent protein domains that bind to specific cellular structures or other proteins during translation elongation. This has been shown for polarity proteins in C. elegans, centrosomal proteins in human cells and D. melanogaster embryos, and a variety of other proteins in human cells (Chouaib et al., 2020; Parker et al., 2020; Safieddine et al., 2021; Tocchini et al., 2021; Winkenbach et al., 2022).

Despite the broad observation of localized translation across organisms, the underlying mechanisms and importance for animal development remain largely unexplored. For example, it is unclear whether mRNA hotspots result from random diffusion or directed movement, and what the time interval is between initiation of translation and localization. Furthermore, it is unknown whether mRNA-ribosome complexes are ‘trapped’ or dynamically move between their hotspots and cytosol, and whether translation always results in mRNA localization or if some translated mRNAs stay in the cytoplasm despite being translated. Understanding the importance of localized translation is further complicated by the fact that translation-dependent mRNA localization is often guided by a localization signal within the protein. Consequently, interference with these signals may affect protein function independent of localized translation. Finally, there is little understanding of when and where proteins are synthesized during development, as there are currently only few assays that visualize translation dynamics in living organisms.

Here, we developed and employed SunTag-based live imaging of translation on single mRNAs in the nematode C. elegans, and examine the dynamics and importance of localized translation. We specifically focus on ERM-1, which is the sole ortholog of the human Ezrin/Radixin/ Moesin (ERM) family and plays a key role in organizing the apical cell cortex and lumen formation in tubular epithelia (Gobel et al., 2004; Van Furden et al., 2004). ERM proteins link phospholipids and proteins embedded in the plasma membrane to the underlying actin cytoskeleton. Loss of *erm-1* results in developmental defects and lethality, underscoring its critical role in animal development (Gobel et al., 2004; Ramalho et al., 2020; Van Furden et al., 2004). Recent research has demonstrated that *erm-1* mRNA co-localizes with ERM-1 protein at the plasma membrane (Li et al., 2021; Parker et al., 2020; Winkenbach et al., 2022). This co-localization is dependent on translation and the presence of nascent polypeptides, as inhibiting translation or mutation of the membrane-binding domain of ERM-1 diminishes *erm-1* mRNA localization (Winkenbach et al., 2022). Based on these observations, localized translation at the membrane and translation-dependent mRNA localization were proposed for *erm-1* (Winkenbach et al., 2022). Here, we apply SunTag-based live imaging and show that most of the endogenous *erm-1* translation indeed occurs at the membrane. We observed that localized *erm-1* translation is highly dynamic and heterogeneous: some translated mRNAs stay in close proximity of the membrane for minutes, while others lose the interaction and move back into the cytoplasm, even in the presence of translation. This highlights the importance of live-tracking of this dynamic process and excludes a simple model where every translated mRNA is trapped at the plasma membrane. In addition, we demonstrate that localized translation is required for proper ERM-1 function. By combining the SunTag translation system with the PP7/PCP mRNA tethering system, we re-localized *erm-1* translation to the nuclear envelope, away from its preferred location at the plasma membrane. This re-localization of *erm-1* affects the ability of ERM-1 to link the actin cytoskeleton with the apical membrane of intestinal cells during development, suggesting that localized translation is important for fast posttranslational processing, modification, or complex formation of ERM-1. These findings underscore the potential of the SunTag system for investigating translation dynamics in animal development and emphasize the importance of localized translation and translation-dependent mRNA localization for protein function and development.

## Results

### Visualizing translation of single mRNAs in C. elegans embryos and larvae

To visualize the translation of single mRNAs during development, we adapted the microscopy-based SunTag system for use in C. elegans. The Suntag live-cell imaging system combines two components that together allow the microscopic detection of nascent polypeptides. For one, a SunTag peptide array (generally 24xGCN4 peptides) is incorporated at the 5’-end of a gene of interest. This is combined with co-expression of the anti-GCN4 single-chain variable antibody fragment (scFv) fused to eGFP (scFv::GFP; SunTag antibody). When the mRNA is translated, the GCN4 peptide array emerges from the ribosome first and is rapidly recognized by the cytoplasmic scFv::GFP antibodies. The binding of multiple antibodies to one nascent protein, combined with translation of a single mRNA by multiple ribosomes leads to amplification of the GFP signal. Thus, the translation of an individual mRNA can be observed as a bright fluorescent signal spot (Morisaki et al., 2016; Pichon et al., 2016; Ruijtenberg et al., 2020; Tanenbaum et al., 2014; Wang et al., 2016; Wu et al., 2016; Yan et al., 2016).

To develop the SunTag system for C. elegans, we first tested if the SunTag antibody is efficiently recruited by a single or few GCN4 SunTag epitopes fused to an endogenous protein (Fig. 1A,B). We expressed a C. elegans optimized version of the scFv::GFP SunTag antibody under the control of the ubiquitous *eft-3* promoter (Aljohani et al., 2020; Tanenbaum et al., 2014). This resulted in uniform scFv::GFP expression throughout the cytoplasm and somewhat higher fluorescence in the nucleus of somatic tissues and the germline (Fig. 1C). We then fused the SunTag epitope(s) to the nuclear pore protein NPP-9, the polarity protein PAR-3, and the membrane-cytoskeleton linker protein ERM-1 to assay antibody recruitment. As expected, we observed clear re-localization of GFP to the nuclear pores, the anterior domain of embryonic blastomeres, and the plasma membrane, respectively (Fig. 1D-F). Membrane enrichment of ERM-1 was comparable between the ERM-1::SunTag and an ERM-1::GFP strain (Fig. 1F, right panel), further confirming that the scFv::GFP antibody can efficiently label GCN4 epitope-tagged proteins in *C. elegans*.

**Figure 1:**
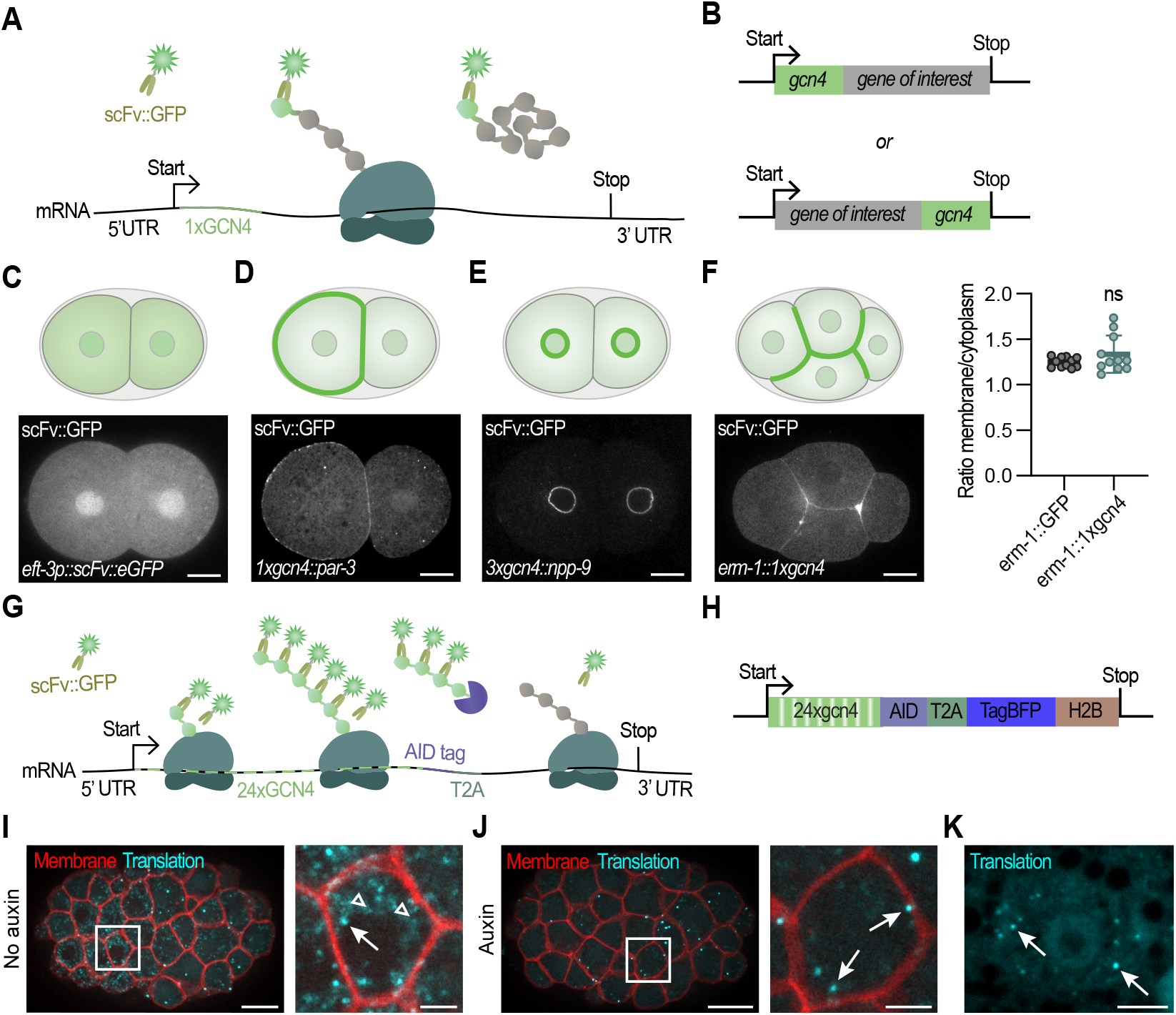
Visualizing proteins and translation of single mRNAs in *C. elegans* embryos and larvae using SunTag. **(A-B)** Schematic overview of the SunTag-based protein labeling strategy. Expression of the protein of interest fused to the GCN4 peptide leads to binding and localization of the SunTag antibody (scFV::GFP), allowing visualization of genes of interest. **(C)** Schematic overview and representative image of a 2-cell stage *C. elegans* embryo expressing the SunTag antibody (scFv::GFP). **(D)** Schematic and representative image of PAR-3 expression using the SunTag system. **(E)** Schematic and representative image NPP-9 expression using the SunTag system. **(F)** Schematic and representative image of ERM-1 expression using the SunTag system (left) and quantification of membrane over cytoplasmic ratio of ERM-1 levels using either conventional GFP labeling of ERM-1 or the SunTag system (right). Each dot represents a single embryo (n=11, 11). Test of significance: Unpaired Student’s T-test with Welch’s correction, with significance set at p≤0.05. ns = not significant **(G)** Schematic of translation imaging strategy using the SunTag system. **(H)** Schematic overview of our translation imaging reporter, which contains 24xGCN4 peptides, an AID sequence, a T2A ribosome skipping site, the coding sequence of BFP::H2B and the tbb-2 3’UTR. Introducing the AID sequence C-terminally of the SunTag peptides, in combination with the T2A-system, allows specific degradation of mature SunTag peptides while leaving translation sites unaffected. **(I-J)** Representative images of an embryo expressing the SunTag antibody (scFv::GFP; depicted in cyan), the translation imaging reporter (*eft-3p::24xGCN4::AID::T2A::BFP::h2b::tbb-2 3’UTR*), the membrane marker (*pie-1p::mCherry::PH*; depicted in red) and TIR protein (*eft-3p::TIR1::F2A::mTagBFP2::AID::NLS*; not shown) without auxin treatment (I) or with auxin treatment (J). Arrows in insets indicate translation spots, open triangles indicate (mature) GCN4 proteins. Scale bar: 10 µm (large panel); 2.5 µm (small panels). **(K)** A representative image of the seam cell plane (skin cells) of an L2 larvae is shown, expressing scFv::GFP and the translation imaging reporter *eft-3p::24xGCN4::AID::T2A::BFP::h2b::tbb-2 3’UTR*. Translation spots in cyan (arrows). Inset to the right. Scale bar: 20 µm (left); 5 µm (zoom-in).

Next, we used the SunTag system to visualize translation. We designed a reporter encoding 24xGCN4 SunTag peptides, followed by BFP::H2B under the control of the *eft-3* promoter. This allowed simultaneous visualization of translation (based on the SunTag peptides) and final protein levels (based on BFP::H2B) (Fig. 1G,H). To ensure that the SunTag does not interfere with BFP::H2B expression, we separated the 24xGCN4 SunTag peptides from the BFP::H2B by using a T2A ribosome skipping site, resulting in two distinct proteins from the same mRNA in approximately a 1:1 ratio (Fig. 1G,H) (Ahier & Jarriault, 2014). Combined expression of the reporter and the SunTag antibody resulted in the appearance of bright green fluorescent translation spots in both embryos and larvae, across different tissues (Fig. 1I, arrows). Control embryos solely expressing the SunTag antibody without the SunTag peptides did not exhibit any translation spots (Fig. 1A). Of note, the SunTag reporter is silenced in early embryos and the first translation spots appear around the 16-cell stage. In addition to the bright translation spots, we observed numerous dim GFP-positive spots (Fig. 1I, open arrowheads). We hypothesized that these dimmer spots are mature proteins consisting of only one SunTag peptide array. Therefore, we combined the SunTag system with the Auxin Inducible Degron (AID) system (Nishimura et al., 2009; Sepers, Verstappen, et al., 2022; Zhang et al., 2015). The AID system enables targeted degradation of AID-tagged proteins upon addition of auxin and expression of TIR-1, a plant factor that in complex with auxin mediates AID recognition by SCF-like E3 ligases. By introducing the AID sequence C-terminally of the SunTag peptides and upstream of the T2A sequence, we expected to specifically degrade the mature SunTag polypeptides without affecting the SunTag signal at the translation sites (Fig. 1G,H)(Wu et al., 2016). Indeed, the dimmer spots largely disappeared upon addition of auxin, indicating that they represent mature proteins rather than translation sites (compare Fig. 1I with Fig. 1J (embryo); Fig. 1K (seam cells)). Thus, by combining the AID system and the SunTag system we can specifically visualize translation with a high sensitivity.

### Validation of the SunTag system

To validate whether the bright GFP spots are active sites of translation, we made two assumptions: 1) the spots should rapidly disappear upon translation termination and 2) they should co-localize with mRNAs. When we inhibited translation, by subjecting the animals to a short heat shock, we observed a rapid and significant reduction in the number of translation spots, in support of our first assumption (Fig. 2A,B). Next, we assessed co-localization between individual mRNAs and presumed translation spots, in both fixed and live embryos. In fixed samples, we combined single-molecule inexpensive fluorescence in situ hybridization (smiFISH) to visualize reporter mRNA with immunofluorescence-based (IF) detection of the scFv::GFP to visualize translation (Fig. 2C,D)(Tocchini et al., 2021). We observed that ∼60% of the GFP clusters coincide with an mRNA spot, supporting our second assumption (Fig. 2D). While most translation spots overlapped with mRNA, a substantial fraction did not. Although some bright GFP spots might be mature proteins or protein aggregates, we expect that this reflects a low labeling efficiency of our mRNA probes (only ∼62%; see methods, Fig. S1A) and consequent failure to detect a fraction of reporter mRNA molecules. Together, this indicates that most or all GFP spots indeed represent active translation. Interestingly, in line with recent literature, we also observed that many mRNAs are not translated (Sonneveld et al., 2020), highlighting the need for single-molecule measurements to explore translation heterogeneity.

**Figure 2:**
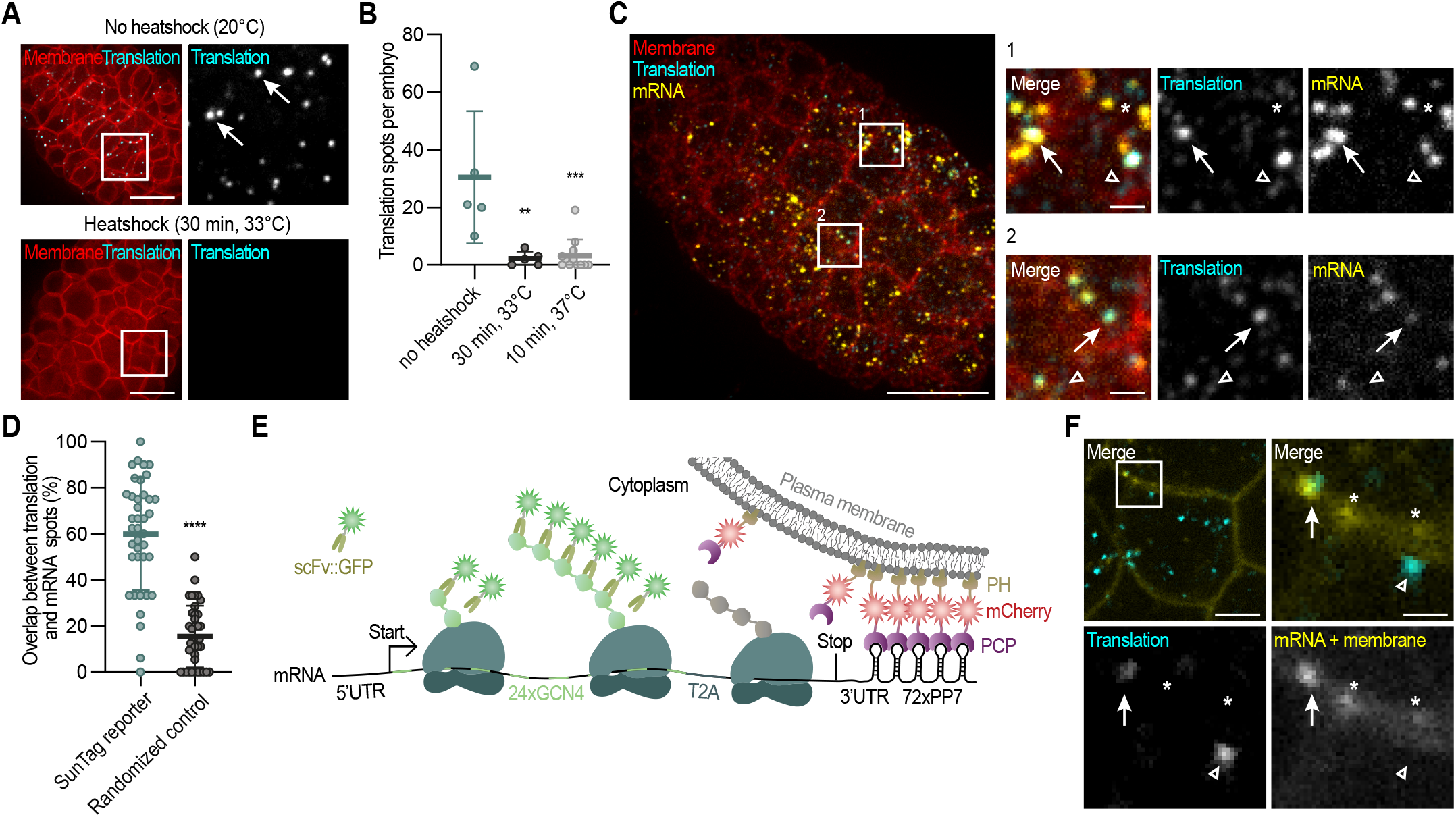
Validations of the SunTag translation imaging system in *C. elegans*. **(A)**Representative images of an embryo expressing the SunTag antibody (scFv::GFP), the translation imaging reporter (*eft-3p::24xGCN4::AID::T2 A::BFP::h2b::tbb-2 3’UTR*), and the membrane marker (*pie-1::mCherry::PH*), subjective to normal culturing conditions (upper panel) or heat-shock (bottom panel). Insets to the right, arrows indicate active translation sites. Scale bar: 10 µm. **(B)** Quantification of the number of translation spots in 30- to 60-cell embryos in non-treated and heat shocked animals. Each dot represents one embryo (n=5, 5, 12), bars display the mean ± SD. Kruskal-Wallis followed by a Dunn’s post-hoc test was used to assess statistical against the “no heat-shock” condition; **p≤0.01 and ***p≤0.001. **(C)** Representative image of an fixed embryo expressing the SunTag antibody (scFv::GFP), a translation imaging reporter (*eft-3p::24xGCN4::AID::T2A::BFP::H2B:20xPP 7:tbb-2 3’UTR*), and a membrane marker (*pie-1p::PCP::mCherry::PH*). Reporter mRNAs (yellow), membranes (red) and translation spots (cyan) were detected using a combination of smiFISH and IF. Insets to the right. Arrows in insets indicate translation spots overlapping with mRNA. Open triangles indicate (mature) GCN4 proteins (weak cyan signal), not overlapping with mRNAs. Asterisks indicates non-translated mRNA. Scale bar: 10 µm (large panel); 1 µm (small panels). **(D)** Quantification of the number of bright cyan spots (presumable translation spots) that overlap with mRNA spots. Each spot represents one embryo (n=39, 39), bars indicate the mean ± SD. Test of significance: Unpaired Student’s T-test with Welch’s correction; ****p≤0.0001. The number of mRNAs may be underestimated due low labelling efficiency (Fig. S1A). **(E)** Schematic overview of the SunTag translation imaging system in combination with mRNA labeling and membrane tethering using the PP7 system. In addition to the reporter described in Figure 1G, 72xPP7 hairpins where inserted into the translation imaging reporter and *pie-1p::PCP::mCherry::PH* was expressed to allow mRNA visualization. **(F)** Representative image of a part of a live embryo, extrachromosomally expressing the reporter as shown in (C), indicating overlap between translation spots (cyan) and mRNAs (yellow). Arrow in insets indicates a translation spot that overlaps with mRNA, open arrowhead indicates a translation spot not overlapping with mRNA and asterisks indicate non-translated mRNAs. Scale bar: 2.5 µm (entire cells); 0.75 µm (zoom-in). See also Fig. S1.

Next, we set out to visualize mRNA and translation simultaneously in live animals. To achieve this, we combined the SunTag system with the PP7/PCP system for mRNA detection (Chao et al., 2008; Li et al., 2021). This involved introducing 72 copies of a PP7 hairpin sequence into the 3’UTR of our reporter gene and co-expressing the PP7 bacteriophage coat protein (PCP) fused to mCherry and a PH domain (PCP::mCherry::PH). The PCP::mCherry::PH proteins localize to the membrane and bind with high affinity to the PP7 hairpin sequences. The 72xPP7 array in the mRNA clusters the PCP::mCherry::PH signal, which allows us to observe mRNAs as individual mCherry-fluorescence spots (Fig. 2E). Using this approach, we often observed the overlap between mRNA and translation spots (Fig. 2F). However, we also observed translation spots that not overlapped with mRNA, which may be due to a low mRNA detection by the PP7/PCP system in *C. elegans* or by low PCP::mCherry::PH levels (see discussion). Interestingly, tracking of membrane-tethered SunTag mRNAs over time revealed the separation of a bright GFP spot that stayed at the membrane and a dim GFP spot that diffused away. This dim spot is most likely an individual, fully synthesized SunTag protein, released from the translation site (bright spot) (Fig. S1B). Together these results show that the bright GFP SunTag spots are sites of active translation, and highlight its powerful potential to study translation dynamics of single mRNAs in real-time in live C. elegans embryos and larvae.

### Localized translation of endogenous *erm-1* mRNA

In the *C. elegans* embryo, localized translation has been suggested for several polarized proteins, including DLG-1, AJM-1 and ERM-1 (Parker et al., 2020; Tocchini & Mango, 2024; Tocchini et al., 2021; Winkenbach et al., 2022). We applied the SunTag system to study the dynamics of localized *erm-1* translation in developing embryos. Similar to our reporter, we inserted 24 SunTag peptides followed by a T2A sequence between the start codon and coding sequence of endogenous *erm-1a* (Fig. 3A), to allow visualization of translation directly from the start of the process. We combined this with expression of the scFv::GFP SunTag antibody and a membrane marker (*pie-1p::mCherry::PH*) (*erm-1*^*24xSunTag*^ strain). The inclusion of the T2A sequence is important as the N-terminal protein fusion of ERM-1 can interfere with its function (Amieva et al., 1999; Henry et al., 1995; Roy et al., 1997). We observed a slightly lower growth rate, but no developmental or morphological defects related to loss of erm-1 in our *erm-1*^*24xSunTag*^ strain. We first examined *erm-1*^*24xSunTag*^ mRNA levels using smiFISH. In contrast to our *erm-1::GFP* control, *erm-1*^*24xSunTag*^ transcripts could not be observed in the early embryos, which is probably due to germline silencing. However, multiple *erm-1*^*24xSunTag*^ transcripts appeared between the 16- and 32-cell stage and expression was abundant during later stages. From the 40-cell stage, there was no difference in either the number or location of *erm-1* transcripts between the *erm-1::GFP* control and *erm-1*^*24xSunTag*^ animals (Fig. S2A). Next, we examined translation of *erm-1*^*24xSunTag*^ using live-imaging. The first translation spots became visible from the 32-cell stage onward, consistent with the observed mRNA expression patterns. Around the 64-cell stage, multiple translation spots emerged at the plasma membrane, across the entire embryo (Fig. 3B). By the bean stage, these translation spots were particularly enriched in the intestinal progenitor cells (Fig. S2B), one of the tissues where ERM-1 is highly expressed and where its function is crucial (Ramalho et al., 2020). Taken together, we created an endogenously SunTag-tagged *erm-1* strain, which allows us to study translation localization of *erm-1a* over the course of development, except for the earliest stages of embryonic development.

**Figure 3:**
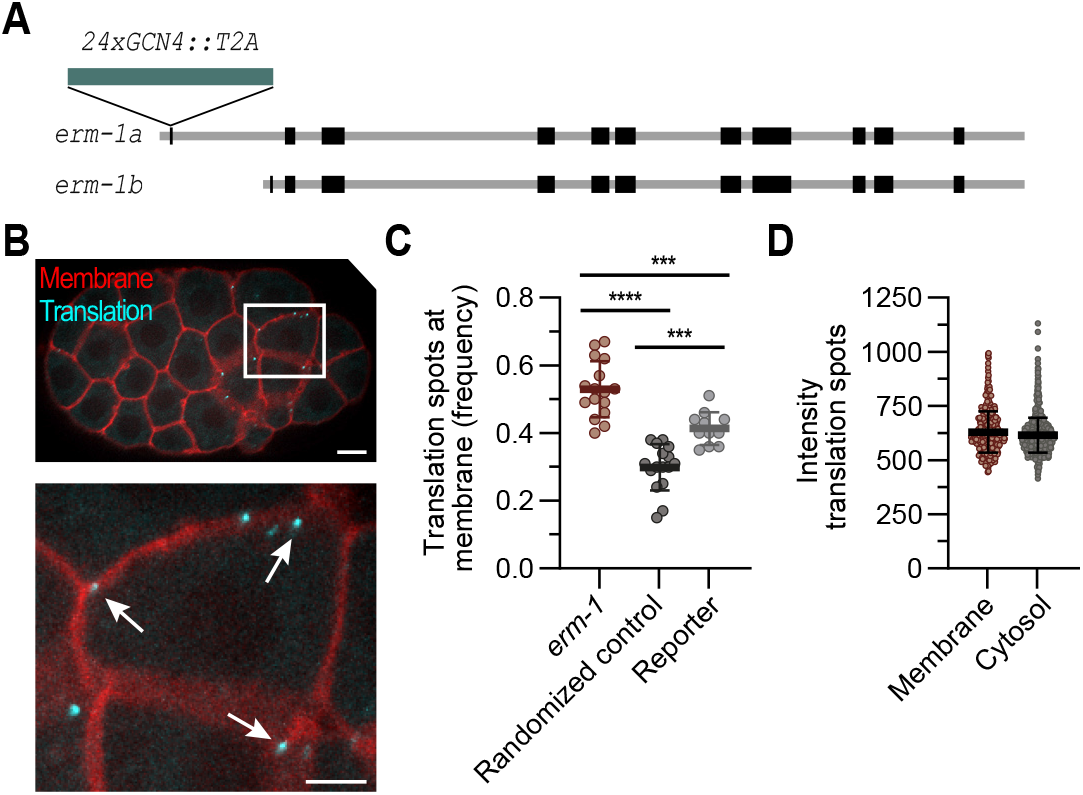
Translation of the *erm-1* is enriched at the plasma membrane. **(A)**Schematic of the *erm-1* locus with the *erm-1a* and *erm-1b* isoforms. To study translation of *erm-1*, 24xGCN4 peptides are inserted directly downstream of the *erm-1a* start codon. **(B)** Image of a representative embryo expressing *24xGCN4::T2A::erm-1*, the SunTag antibody (scFv::GFP) and the membrane marker (mCherry::PH). Membranes in red, translation sites in cyan. Inset shown at the bottom. Arrows indicate translation sites in close vicinity of the membrane. Scale bar: 10 µm (embryo); 5 µm (zoom-in). **(C)** Quantification of the number of translation spots within 0.35 µm of the membrane in the *24xGCN4::T2A::erm-1* strain, in a randomized computed control and in the *eft-3p::24xGCN4 ::AID::T2A::BFP::h2b::tbb-2 3’UTR* strain. Each dot represents the percentage of spots in the proximity of the membrane in a single embryo (n=16, 16, 11). Test of significance: One-way ANOVA with Bonferroni’s post-hoc test. ***p≤0.001 and ****p≤0.0001. **(D)** Quantification of the intensity of translation spots at the membrane and in the cytosol. Each dot represents a single translation spot (n=480, 1744). See also Fig. S2.

To determine whether translation is indeed enriched at the membrane, we quantified the percentage of spots that resided within 0-0.35 micron distance from the membrane. We found an average of 53% (ranging from 40-67% in individual embryos) of translation spots localized at the membrane, which is significantly more than either our SunTag BFP::H2B reporter (41% resided within 0-0.35 range) or a computed Z-flipped control (30% of the spots resided within 0-0.35 range) (Fig. 3C). No difference in spot intensity was observed between *erm-1* translation at the membrane or in the cytoplasm (Fig. 3D). Noteworthy, the 54% enrichment in *erm-1* translation at the membrane is higher than the 40% membrane enrichment of *erm-1* mRNA observed by the Nishimura lab, which fits the model that translated mRNAs have an increased chance of being located close to the membrane (Winkenbach et al., 2022). In *C. elegans* embryonic blastomeres, the ERM-1 protein localizes to the entire plasma membrane. However, in mid-embryonic stages, when intestinal cells undergo polarization and the apical domain is formed, ERM-1 protein becomes almost exclusively localized to the apical membrane (Gobel et al., 2004; Van Furden et al., 2004). This apical localization of ERM-1 is hypothesized to occur in a two-step process, where ERM-1 proteins first localize to membranes in general (because of FERM domain interactions with PIP_2_ in the membrane) after which phosphorylation of ERM-1 leads to the specific apical localization (Fievet et al., 2004; Ramalho et al., 2020; Yonemura et al., 2002). Based on mRNA localization patterns in control and ERM-1 phosphorylation-defective mutants, it has been suggested that the movement of ERM-1 protein from the basolateral to the apical membranes occurs post-translationally (Winkenbach et al., 2022). Indeed, in mid-stage embryos there is an enrichment of translation at both the lateral and apical membrane (Fig. S2B). This demonstrates that *erm-1* is translated near the membrane, but not exclusively at the apical site, where ERM-1 function is required for apical membrane morphogenesis and tube formation.

### Dynamics of localized translation

The current model of localized *erm-1* translation (Li et al., 2021; Parker et al., 2020; Winkenbach et al., 2022) proposes that *erm-1* mRNAs rapidly locate to the membrane upon translation initiation. However, this is a hypothesis as visualization of *erm-1* translation was not feasible before, and the dynamics and heterogeneity of translated mRNAs are unknown. Using timelapse imaging and single-molecule tracking of *erm-1* translation spots, we identified a variety of different mRNA translation behaviors (Fig. 4A). First, we observed *erm-1* translation events that started in the cytoplasm, and over time localized to the membrane, where they then remained for the duration of the recording (5 minutes) (Fig. 4A, arrows; Fig. 4B), in line with the proposed model. Second, we found translated mRNAs that moved throughout the cytoplasm without interacting or staying close to the membrane, indicating that that localization can take longer than 5 minutes or that translation does not always lead to localization to the membrane (Fig. 4A, open arrowhead). Third, we observed mRNAs that where already near the membrane at the start of the recording and stayed there for the entire 5-minute duration (Fig. 4A, asterisks; Fig. 4C). Despite their localization near the membrane, these mRNAs were highly mobile, moving laterally across the membrane. This lateral movement may be due to membrane fluidity or constant interaction and dissociation between PIP_2_ and ERM-1 Furthermore, this could indicate multiple rounds of translation where one nascent protein is released before the next nascent FERM domain binds PIP_2_. During this brief interval, the mRNA may move freely near the membrane, before interacting with PIP_2_ again. As such, translated mRNAs might continuously shuttle between the cytoplasm and the area in the membrane’s vicinity.

**Figure 4:**
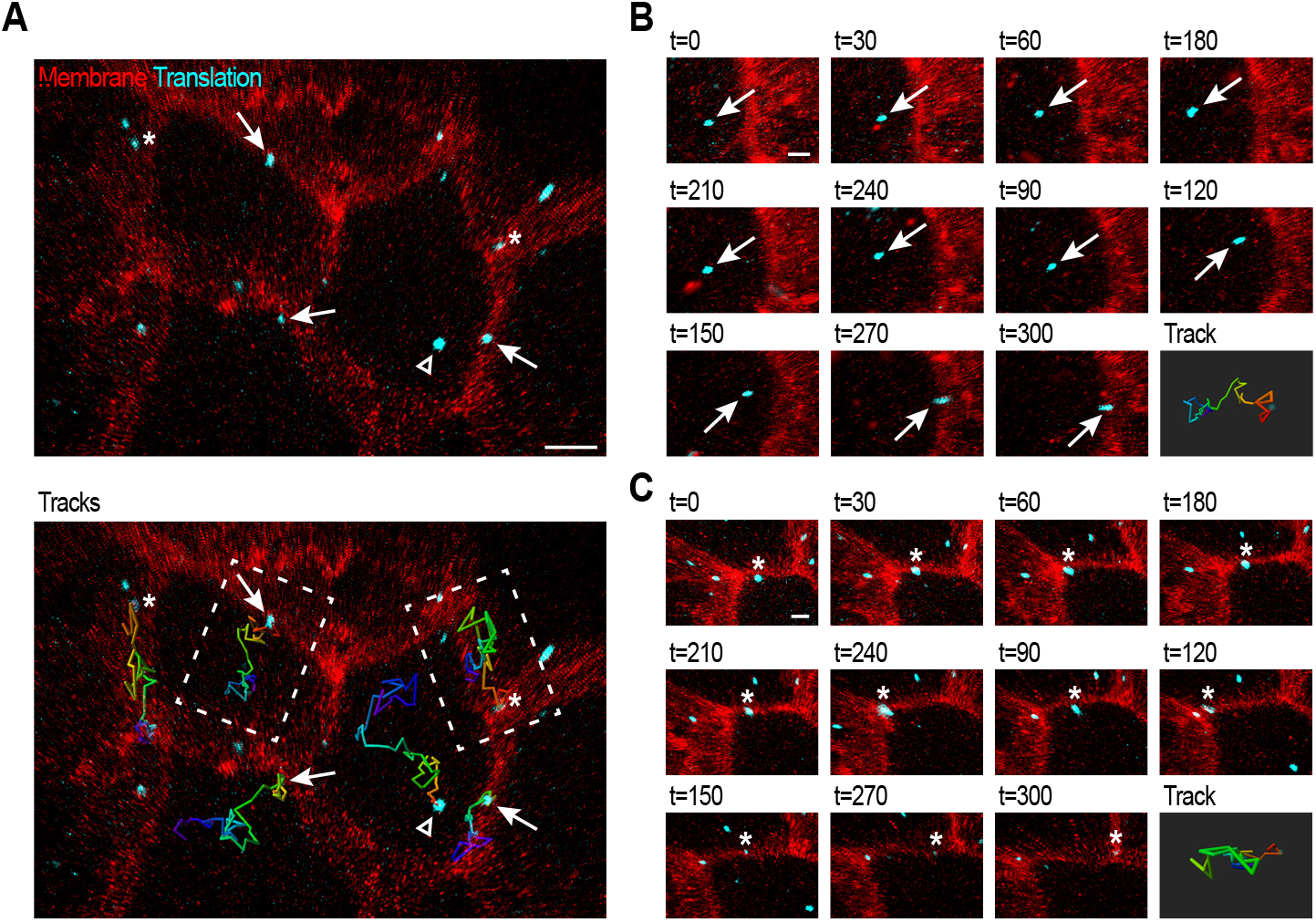
*erm-1* translation spots show different dynamic behaviors. **(A)**Live tracking of individual translation sites in animals expressing *24xGCN4::T2A::erm-1*, the SunTag antibody (scFv::GFP), and the membrane marker (mCherry::PH). Translation spots are shown in cyan, membranes in red. Bottom panel shows the movement-tracks of individual spots over time. Dark blue represents the start of the track, red the end of the track. Arrows indicate translation spots that start in the cytoplasm and localize to the membrane. The open triangle indicates a translation spot that starts and stays in the cytoplasm over the course of the movie. Asterisks indicate translation spots that are located in proximity of the membrane from start to finish of the recording. **(B)** Time series of left inset showing the localization of an individual translation spot to the membrane over the course of 300 seconds. **(C)** Time series of right inset showing the movement of an individual translation spot in close proximity of the membrane for 300 seconds. Scale bar: 2.5 µm (embryo); 1 µm (zoom-in).

Movement towards the membrane could either be directed by active transport or dependent on random diffusion followed by anchoring at the membrane. Our tracks show signs of both non-directed movements (for example blue part, mRNA track, Fig. 4B), as well as rapid, directional movement (for example green part, mRNA track, Fig. 4B). These switches between movement categories have been reported for *erm-1* mRNA before, by using live imaging of *erm-1* reporter mRNAs based on the PP7-PCP system, and were attributed to transitions between a ribosome-free and a ribosome-bound state (Li et al., 2021). However, our data indicates that also in the presence of translating ribosomes, the speed and directionality of mRNA movement changes over time.

### Localized translation of *erm-1* is important for ERM-1 function

Localized translation of *erm-1* could be functionally important, for example, to regulate ERM-1 expression, localization, or to ensure fast post-translational modification of ERM-1. Alternatively, localized translation may not have a specific function and could merely be a byproduct of co-translational binding of nascent ERM-1 proteins to the plasma membrane via the PIP_2_ binding domain, which could drag the whole mRNA-ribosome complex along. It has been demonstrated that altering ERM-1 protein localization or function affects *erm-1* mRNA localization at the membrane (Winkenbach et al., 2022). Here, we pose the reverse question: if we alter the cellular site where ERM-1 is synthesized, does this impact ERM-1 protein localization and/or function? For this purpose, we re-localized *erm-1* mRNAs to nuclear pores, relatively distant from the plasma membrane. We used the PP7/PCP mRNA tethering system, and expressed PCP::mCherry fused to the endogenous nuclear pore protein NPP-9 (PCP::mCherry::NPP-9) (Fig. 5A,B). We first tested if reporter mRNAs could be re-localized to the nuclear pores, by incorporating 20xPP7 hairpins in the SunTag::BFP::H2B control reporter. This led to a clear re-localization of the SunTag spots, without causing developmental defects (Fig. S3A). Thus, our system efficiently re-localizes mRNAs, and enhancing translation at the nuclear pore has no detrimental effects for animal development.

**Figure 5:**
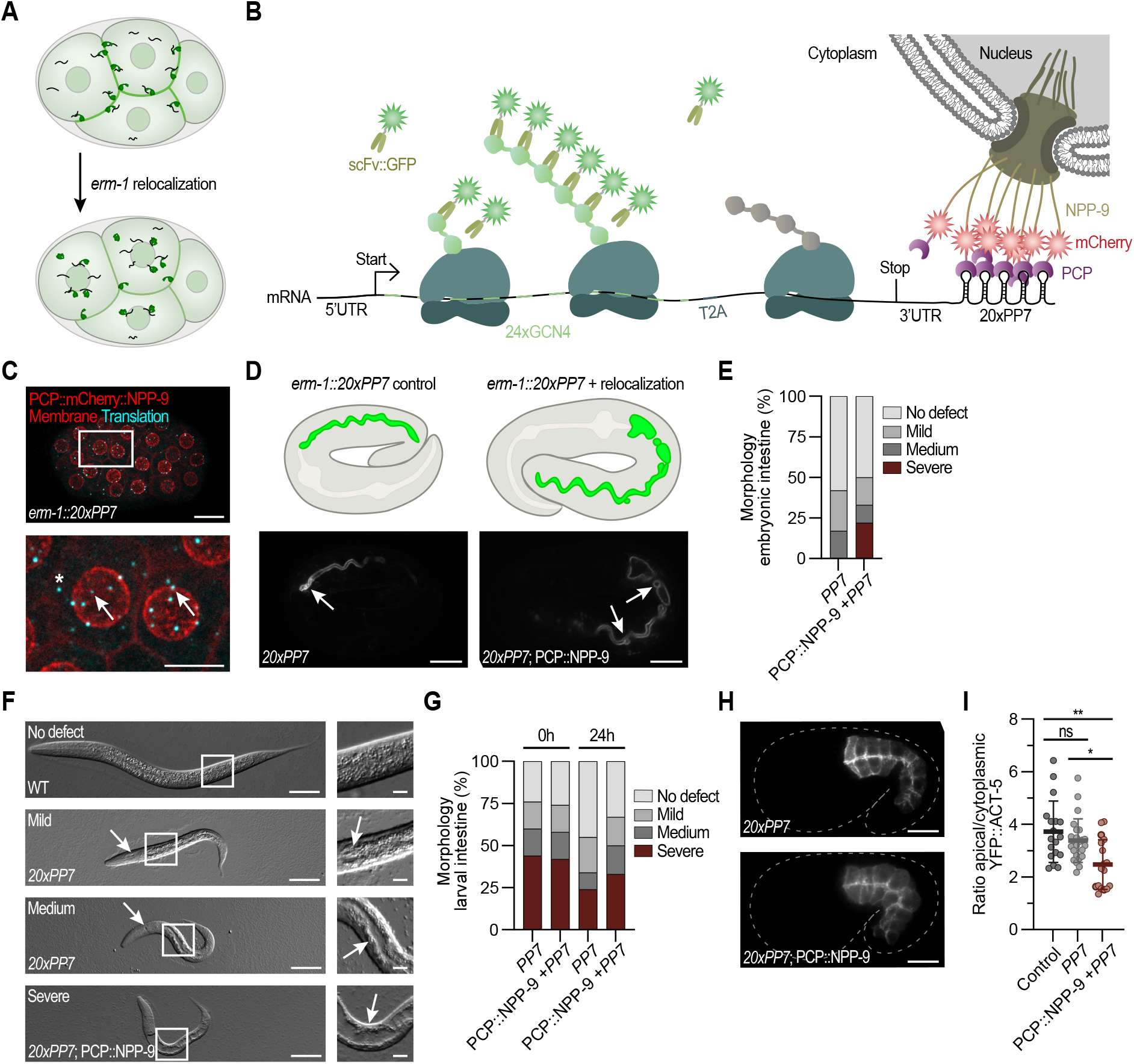
Disruption of localized *erm-1* translation affects ERM-1 protein function. **(A)** Schematic overview of the experimental design to re-localize *erm-1* mRNAs from the membrane to the nuclear pore to address the importance of translation in close proximity of the membrane. **(B)** Schematic overview of SunTag-based translation imaging of *erm-1* in combination with mRNA re-localization and tethering to the nuclear pore, using the PP7 system. For this, 20xPP7 sites are inserted in the 3’UTR of erm-1 and PCP::mCherry is expressed from the endogenous locus of NPP-9. **(C)** Representative image of an embryo expressing *24xGCN4::T2A::erm-1::20xPP7*, PCP::mCherry::NPP-9 (red), mCherry::PH (red). The majority of the translation spots (cyan) can be observed around the nucleus. Inset below. Arrows indicate spots at the nucleus. Asterisks indicates a cytoplasmic spot. Scale bar: Scale bar: 10 µm (embryo); 5 µm (zoom-in). Quantification of the re-localization efficiency can be found in Fig. S3B. **(D)** Representative image of embryos expressing ACT-5::GFP. Arrows indicate intestinal defects (constrictions and widened lumen) upon introduction of *20xPP7* sites in the 3’UTR of *erm-1* (left) and *erm-1* mRNA re-localization (right). Scale bar: 10 µm. **(E)** Quantification of morphological defects in embryos with or without *erm-1* re-localization (n=12, 36). **(F)** Representative DIC images, showing the larval defects observed in the indicated genotypes. Insets shown to the right, arrows indicate morphological defects in the intestine. Scale bar: Scale bar: 50 µm (larvae); 10 µm (zoom-in). **(G)** Quantification of morphological defects in larvae with or without *erm-1* re-localization, 0 and 24 hours after hatching (n=25, 19, 29, 18). **(H)** Intestinal distribution of YFP::ACT-5 in embryos with (upper panel) or without (lower panel) *erm-1* re-localization. Scale bar: 10 µm. **(I)** Quantification of the apical to cytoplasmic ratio of YFP:ACT-5 levels. Each dot represents one embryo (n=18, 24, 17). Error bars indicate mean ± SD. Test of significance: Kruskal-Wallis with Dunn’s post-hoc test. ns = not significant, *p≤0.05, **p≤0.01. See also Fig. S3.

We next applied our tethering technique to *erm-1* by introducing 20xPP7 hairpins in the 3’UTR of *erm-1*, resulting in the *erm-1*^*24xSunTag;20xPP7*^ strain. As expected, when combining PCP::mCherry::NPP-9 expression with *erm-1*^*24xSunTag;20xPP7*^ expression, we observed that the majority (82%) of the translation spots now resided at the nuclear envelop, instead of the plasma membrane (only 12%, compared to 53% in non-re-localized strains) (Fig. 5C, Fig. S3B). Interestingly, these animals displayed severe developmental defects characteristic of *erm-1* mutants, such as cysts in the intestinal lumen and excretory canal in both embryos and larvae (Fig. 5D-G). Specifically, 22% of embryos showed severe morphological changes of the intestinal lumen (Fig. 5E) which increased to 47% in L1 larvae (Fig. 5G). Previous research showed that *erm-1* phosphorylation mutants can recover from these defects over time (Ramalho et al., 2020), and indeed, only 28% of the animals showed severe developmental defects 24 hours later (Fig. 5G).

In a control *erm-1*^*24xSunTag;20xPP7*^ strain, containing the 20xPP7 hairpins in *erm-1* but not expressing PCP::mCherry::NPP-9, no re-localization of *erm-1* translation was observed. However, this strain also showed defects in lumen formation, although less severe compared to the re-localization strain (Fig. 5E,G). For example, no severe morphological defects were observed in embryos (Fig. 5E), and in larvae the percentage of animals with severe defects was slightly lower compared to the *erm-1* re-localization strain (Fig. 5G). IF-based detection of ERM-1 showed that introduction of the 20xPP7 hairpins in the 3’UTR led to an overall reduction of ERM-1 expression, regardless of whether the mRNAs were re-localized or not (Fig. S3C; due to technical variation, small differences may have been missed). The lower ERM-1 expression could be caused by the increased *erm-1*^*24xSunTag;20xPP7*^ 3’UTR length, which may sensitize mRNAs for degradation by the nonsense mediated decay machinery (Tocchini & Mango, 2024). In line with this, both the re-localization strain and the control strain showed some general developmental defects such as an increased percentage of embryonic lethality and slower growth rates compared to wild-type animals. However, in contrast to the *erm-1* specific phenotypes, there was no significant difference between the two strains in these more general phenotypes (Fig. S3D,E).

Finally, we aimed to address the downstream consequences of the disturbed *erm-1* translation patterns. Accurate ERM-1 expression and phosphorylation is crucial for its role in linking the actin cytoskeleton to the apical membrane of epithelial cells (Hipfner et al., 2004; Ramalho et al., 2020). The apical actin network in *C. elegans* intestinal cells mainly comprises of the specialized actin ACT-5 (MacQueen et al., 2005). The integrated transgene YFP::ACT-5 was used previously to assess actin distribution in *erm-1* mutant animals. Here, loss of ERM-1 function resulted in the loss of YFP::ACT-5 from the apical membrane, accompanied by elevated levels of cytoplasmic and basolateral actin (Ramalho et al., 2020; Sepers, Ramalho, et al., 2022). Therefore, we assessed if YFP::ACT-5 expression is affected in nuclear pore re-localized *erm-1* and non-localized *erm-1* control strains. As expected, we detected a strong ACT-5 enrichment at the apical domain of intestinal cells in wild-type mid-stage embryos. In contrast, in our *erm-1* re-localization strain, apical enrichment of ACT-5 was diminished, whereas ACT-5 concentration was elevated in the cytoplasm and along the basolateral membrane (Fig. 5H,I). These data suggest that localized translation of ERM-1 is important for the immediate recruitment of actin to the apical membrane. Importantly, the reduced apical/cytoplasmic ratio is dependent of *erm-1* mRNA re-localization as the ratio is unchanged in the control *erm-1*^*24xSunTag;20xPP7*^ strain (Fig. 5I). Thus, while introduction of 20xPP7 hairpins into the *erm-1* 3’UTR leads to reduced expression levels and morphological and developmental defects, the remaining ERM-1 is capable of accurately localizing ACT-5. In contrast, ERM-1 protein produced at the nuclear pore has, at least partially, lost this ability, suggesting that the cellular site of translation contributes to ERM-1 function.

## Discussion

In this study, we report the development, validation and use of the SunTag-based single-molecule translation imaging system in *C. elegans*. Using our system, we followed translation of individual mRNAs live and detected localized translation for the membrane-cytoskeleton linker protein ERM-1. Our data indicate that localized translation of *erm-1* at the membrane is important for ERM-1 function in recruiting actin to the apical membrane, and thus for animal development and tissue morphology.

### The SunTag system as a general tool to study translation and visualize proteins of interest

Using the SunTag system we obtained unique insights in the translation dynamics of *erm-1*. In addition, we envision that the SunTag system has a broad application to many more aspects of *C. elegans* development in which the localization and efficiency of mRNA translation is important, including neuronal development, cell cycle regulation and early embryogenesis (Das et al., 2021). In the early *C. elegans* embryo, transcription is silenced, and the control of accurate protein levels thus fully relies on posttranscriptional regulation of gene expression, including translation (Seydoux & Fire, 1994). Being able to visualize the translation dynamics will shed new light on how translation control contributes to cell fate specification and development. Beyond visualizing translation, the SunTag system can be used to visualize and study proteins of interest. Due to its small size of only 19 amino acids, multiple SunTag peptides can easily be fused to genes of interest for amplification of a fluorescent detection signal. Fluorescence amplification can be of great benefit for single molecule imaging (Hu et al., 2023) or visualization of lowly expressed proteins. Moreover, the SunTag system can be used to visualize and measure expression of highly dynamic proteins (e.g., transcription factors), which are challenging to observe using conventional fluorophores because of long maturation times (Bothma et al., 2018; Chang & Dickinson, 2022; Tanenbaum et al., 2014). The SunTag system overcomes this issue as the pre-maturated SunTag antibodies immediately visualize proteins that are fused to the SunTag peptides (Dufourt et al., 2021; Vinter et al., 2021). Taken together, our SunTag system allows live, single molecule imaging of translation in *C. elegans* and provides a framework for understanding the role and underlying mechanisms of translation regulation in *C. elegans* development.

### Considerations when applying the SunTag system and PP7/PCP system in C. elegans

Visualizing mRNAs and their translation provides valuable quantitative information on posttranscriptional gene regulation. However, both the PP7/PCP and SunTag-systems require genetic manipulation that can potentially affect expression of the target gene. For example, we noticed that integrating the SunTag array into the endogenous locus of *erm-1* resulted in germling silencing. Reducing the number of SunTags, removing piRNA sites from the SunTag sequences (Wu et al., 2018; Zhang et al., 2018) or introducing *smu-1* introns (Aljohani et al., 2020) may help overcome germline silencing in the future. Similarly, we found that introducing PP7 hairpins in the 3’UTR of *erm-1* reduced ERM-1 expression (Fig. S3C), probably because longer UTRs are more prone to be recognized and degraded by the nonsense mediated decay machinery (Buhler et al., 2006; Eberle et al., 2008; Hogg & Goff, 2010). Changing the number or location of hairpins could reduce this effect, as demonstrated previously for *dlg-1* (Tocchini & Mango, 2024). It is important to note that the implications of genetic modifications may vary depending on the gene and experimental conditions. For instance, we noticed that using the same PP7/PCP system to re-localize mRNAs to the plasma membrane (PCP::mCherry::PH; Fig. 2F, and Fig. S1B) was less efficient than to re-localize to the nuclear pore (PCP::mCherry::NPP-9; Fig. 5C, and Fig. S3A,B). This could suggest that NPP-9 proteins allow a more stable tethering compared to PH domains. However, we also found that mRNAs tethered to the plasma membrane remained there for extended periods (Fig. S1B), indicating that PCP::mCherry::PH provided stable tethering. We speculate that PP7-tagged mRNAs are immediately bound by PCP::mCherry::NPP-9 proteins when they exit the nucleus, whereas binding to PCP::mCherry::PH proteins at the plasma membrane may take longer or may be more challenging. Although tethering mRNAs was efficient, visualizing mRNAs using the PP7/PCP system was more demanding. Inefficient mRNA detection could either be because of low PCP::mCherry::PH levels, or because labeled mRNAs were masked by high levels of the co-expressed mCherry::PH. In line with this, we noticed that in older embryos – with lower fluorescent signal at the plasma membranes – mRNAs could more easily be detected (data not shown). Finally, when using the SunTag system to visualize translation in *C. elegans* it is important to note that the scFv::GFP SunTag antibody has some affinity to structures close to epithelial apical junctions (Hu et al., 2023). Furthermore, the mature GCN4 proteins have affinity for structures near the DNA of mitotic cells (data not shown), but this is prevented by fast degradation of GCN4 proteins using the AID system. In conclusion, the SunTag system is a powerful method to visualize and measure translation dynamics in *C. elegans* and specific optimization could further enhance its utility in studying translation across all tissues and developmental stages.

### The importance of *erm-1* translation at the membrane for ERM-1 function

In other species, localized translation has been suggested to be important in either creating local hotspots of protein concentration or for the maturation, processing and posttranslational modification of newly synthesized proteins (Das et al., 2021; Kim et al., 2013). Our data indicate that while localized translation of *erm-1* may not be essential for proper protein localization – though technical variation in immunostaining may have hindered detection of small differences in ERM-1 levels and localization – it could play a role in protein interactions or posttranslational processing and modification of ERM-1. Our *erm-1* re-localization strain showed defects similar to those observed in ERM-1 phosphorylation-mutants, such as reduced apical enrichment of ACT-5. Phosphorylation of ERM-1 generally occurs after membrane binding and is important for maintaining an open conformation that enables ERM-1 to link the membrane and actin cytoskeleton (Fievet et al., 2004; Ramalho et al., 2020; Yonemura et al., 2002). We speculate that *erm-1* translation at the membrane contributes to fast phosphorylation of ERM-1, to produce ERM-1 proteins in the ‘ON’ state, ready to perform their function. This can occur either because ERM-1 remains in an open conformation by immediate membrane binding during translation (Winkenbach et al., 2022), or by co-enrichment of the ERM-1 kinase at the membrane. However, the specific kinase responsible for ERM-1 phosphorylation in C. elegans remains unidentified at this stage, leaving the question of which kinase requires *erm-1* to be translated near the membrane for efficient phosphorylation open for investigation. In case *erm-1* is not translated close to the membrane, as in our *erm-1* re-localization strains, the N-terminal FERM domain and C-terminal C-ERMAD domain of ERM-1 probably form an intermolecular interaction, promoting an inhibited state that remains cytoplasmic and unable to function (Gary & Bretscher, 1995; Li et al., 2007; Magendantz et al., 1995; Pearson et al., 2000), which potentially leads to reduced ACT-5 levels at the apical membrane.

While most *erm-1* mRNAs are translated at the membrane, a substantial fraction is translated in the cytoplasm, raising the question of why translation at the membrane is not important for these mRNAs. The answer may lie in the observation that during early embryogenesis the concentration of ERM-1 protein at the membrane is only ∼1.3 times higher compared to the cytoplasm (based on ERM-1::GFP and ERM-1::GCN4, Fig. 1F). This could mean that during these stages there is an excess of ERM-1 (i.e. not all ERM-1 needs to be in ‘ON’ state) or that phosphorylation of ERM-1 is not yet important, consistent with the absence of morphological defects in *erm-1* (phosphorylation) mutants in early embryos (Ramalho et al., 2020). Localized translation could become essential during cellular differentiation and polarization of the intestinal cells around the bean stage, when ERM-1 starts to localize primarily to the apical membrane (Gobel et al., 2004; Van Furden et al., 2004). At this stage, localized translation would ensure fast activation and function of ERM-1, supporting its specialized role at the expanding apical membrane surface of the intestinal epithelium. While some ERM-1 proteins produced in the cytoplasm may eventually be phosphorylated, this process might be too slow to meet the demands of the rapidly developing embryo.

### Heterogeneity in *erm-1* translation

Even though all ERM-1 proteins contain a PIP_2_ binding domain, not all translated mRNAs are localized to the membrane. Why do some mRNAs localize to the membrane, and other mRNAs not? The first reason might be due to technical limitations. Visualization of translation-dependent mRNA localization requires at least two simultaneously translating ribosomes per transcript. This is because of the insertion of the T2A sequence, which splits the SunTag peptides encoded at the 5’-end (required for visualization) from the ERM-1 PIP_2_ binding domain encoded more downstream (required for localization to the membrane). Therefore, if only downstream ribosomes are present, mRNAs may localize to the membrane, but this would not be detected with our methods. Noteworthy, we did not observe disappearance of signal from mRNAs that did localize to the membrane within our 5-minute timelapse imaging, indicating that translation was efficient enough to maintain at least two ribosomes on the transcript in these cases. Conversely, if only ribosomes at the SunTag peptide sequence are present, we may observe translation without localization. Although some mRNAs might indeed be in early translation stages and localization may have not been initiated yet, we estimate that our 5-minute timelapse duration should be long enough for ribosomes to reach the *erm-1* coding sequence. As ribosomes translocate at speeds ranging from 3 to 18 codons per second (Dao Duc & Song, 2018; Ingolia et al., 2011; Morisaki et al., 2016; Pichon et al., 2016; Wang et al., 2016; Wu et al., 2016; Yan et al., 2016), and because the PIP_2_ binding domain is ∼1000 codons from the start, most ribosomes probably reach the PIP_2_ binding domain within 5 minutes after the first appearance of GFP signal. Taken together, technical limitations may cause an underestimation of the proportion of mRNAs that localize to the membrane, but this is unlikely the sole explanation for the presence of cytoplasmic translation spots. Thus, there probably is biological heterogeneity that causes some translated mRNAs to localize to the membrane but not all.

The translated *erm-1* mRNAs that are not localized at the membrane may either fail to be transported, or do not stabilize at the membrane. Our data indicate that most *erm-1* mRNAs remain at the membrane during translation (Fig. 4A,C), suggesting that de-stabilization is not the primary cause of cytoplasmic translation. Localization of *erm-1* mRNAs to the membrane is thought to be mediated by dynein in a translation dependent manner (Li et al., 2021). One explanation for the observed heterogeneity in translation localization is that only the most efficiently translated mRNAs, with many nascent ERM-1 domains, are transported to the membrane. However, this seems unlikely, as we observed no difference in SunTag signal intensity between cytosolic and membrane-bound translation spots (Fig. 3D). Another explanation is that the recognition or translocation of the *erm-1* mRNA-ribosome complexes is a time-consuming process, as reflected in our timelapse imaging, which shows that localization to the membrane can take several minutes. The mechanisms by which the *erm-1* mRNA-ribosome complexes are recognized for transport and the required adapters remain unknown. Additionally, various biological factors may determine which translated mRNAs are targeted to the membrane and which are not. This raises important questions for future research: What causes the heterogeneity in membrane localization, and why are some mRNAs recognized and transported to the membrane while others are not, despite both undergoing active translation.

## Acknowledgments

We thank S. van Dreumel, A. Scheper and G. Nowee for help with experiments, and N. Boster, J. Waterlander, L. Tilburg and M. Schroeder for help with data analysis. We would also like to thank the Ruijtenberg lab members, S. van den Heuvel, M. Boxem and S. Suijkerbuijk for helpful discussions, and S. van den Heuvel, M. Boxem and S. Suijkerbuijk for critical reading of the manuscript. We thank S. van den Heuvel for support with funding. This work utilized resources from Utrecht University and was financially supported by an NWO-XL grant (OCENW. GROOT.2019.017). We also thank WormBase and the Biology Imaging Center Faculty of Sciences of the Department of Biology at Utrecht University. Some strains were provided by the Caenorhabditis Genetics Center, which is funded by the US National Institutes of Health (NIH) Office of Research Infrastructure Programs (P40 OD010440). We thank M. Boxem for the BOX213 strain, M. Tanenbaum for 24xGCN4 and 72xPP7 containing plasmids and M. Galli for the PCP plasmid.

## Author contributions

Conceptualization: E.v.d.S., S.R, Methodology: E.v.d.S., E.K., E.v.d.M.; Writing – original draft: E.v.d.S., S.R.; Writing – review and editing: E.v.d.S., S.R.; Visualization: E.v.d.S, E.K., M.E., S.R.; Formal analysis: E.v.d.S., E.K.; Supervision: S.R.; Project administration: S.R.

The authors declare no competing or financial interests.

## Data availability

Data are available upon request.

## Supplemental information

Document S1: Figures S1-S3 and Table S3

Table S1. Excel file containing additional data too large to fit in a PDF, related to Methods

Table S2. Excel file containing additional data too large to fit in a PDF, related to Methods

## Methods

### C. elegans strain generation

Details on the generation of individual strains, derived from various backgrounds and approaches are provided in Supplementary Table 2. For stable single copy integration of the SunTag antibody and reporter at MosSCI loci, a plasmid-based CRISPR/Cas9 method was used (Friedland et al., 2013). Injection mixes contained *eft-3p::Cas9* plasmid (50 ng/μL; Addgene #46168), single guide RNA (sgRNA) plasmid (50 ng/μL), repair template plasmid (50 ng/μL) and co-injection marker plasmid (2.5 ng/μL) (used sgRNAs and repair templates are listed in Supplementary Table 2). Offspring expressing the co-injection marker were selected, lysed and genotyped to identify genome-edited worms. For integrations at endogenous loci or SunTag reporter alleles, a Cas9 ribonucleoprotein-based approach was used (Ghanta et al., 2021). Injection mixes contained Cas9 protein (0.25 μg/μL) (IDT), TracRNA (0.1 μg/μL) (IDT), *pRF4 (rol-6(su1006))* (40 ng/μL) co-injection marker (Mello et al., 1991), locus specific crRNA (56 ng/μL) (IDT), and either ssODN (0.11 μg/μL) (IDT) or melted dsDNA (25 ng/μL) repair template (crRNAs and repair templates are listed in Supplementary Table 2). dsDNA repair templates were amplified by PCR using primers with containing 5′ SP9 modifications (IDT) and purified using the NucleoSpin Gel and PCR Clean-up Kit (Macherey-Nagel, Cat. No. 740609.250) prior to injections. For extrachromosomal array generation, animals were microinjected with λ-DNA (50 ng/μl) (Thermo Fisher), repair template plasmid (20 ng/μl) and co-injection marker (2.5 ng/μl) (used plasmids sequences are listed in Supplementary Table 2). Progeny expressing the co-injection marker were selected. Correct genome editing or presence of extrachromosomal arrays was confirmed by PCR and Sanger sequencing (Macrogen Europe). For strains generated by crossing, males were obtained by culturing of L4 hermaphrodites at 30°C for 6 hours, followed by incubation at 20°C for 3 days. These males were backcrossed with hermaphrodites of the same strain and maintained according to standard procedures, before crossing with other strains.

### Molecular cloning

The scFv::GFP was based on the *pHR-scFv-GCN4-sfGFP-GB1-dWPRE* plasmid (Addgene #60907)(Tanenbaum et al., 2014) and germline optimized (Aljohani et al., 2020). To create the repair template plasmids for the SunTag reporter integrations, 24xGCN4 sequences were derived from plasmid *pcDNA4TO-24xGCN4_v4-kif18b-24xPP7* (Addgene #74928)(Yan et al., 2016). PP7 hairpin sequences were a kind gift from the Tanenbaum lab (Hubrecht Institute, Utrecht) and the PCP coding sequence was a kind gift from the Galli Lab (Hubrecht Institute, Utrecht). Primers and G-blocks used for molecular cloning were ordered from IDT. Vectors and DNA fragments were either digested by restriction enzymes (Thermo Scientific or NEB) and ligated by T4 ligase (NEB M0202L) or plasmids were cloned using the Gibson assembly (NEB E2611) cloning strategy (Gibson et al., 2009).

### Auxin mediated degradation of mature GCN4 proteins

For degradation of mature 24xGCN4 proteins in embryos, L4 or young adult animals were transferred to NMG plates containing 1mM auxin (IAA; Alfa Aesar) and seeded with OP50 E. coli, and were incubated overnight. Adult animals were splayed to collect embryos for microscopy.

### Heat-shock experiments

Heat-shock experiments were performed by transferring adult worms to pre-warmed M9. Animals were incubated at 20°C or 33°C for 30 min or at 37°C for 10 min and splayed immediately afterwards to obtain the embryos for microscopy.

### smiFISH

Custom smiFISH primary probes were designed against the *24xGCN4, tagBFP2* or *gfp* mRNA sequences using the Oligostan script in RStudio as previously described (Tsanov et al., 2016). A pool of 21-33 primary probes (IDT) were combined in IDTE buffer (pH 8.0) to a final concentration of 0.833 µM per probe. Primary probe sequences can be found in Supplementary Table 3. Secondary (FLAP-Y) probes labeled with CAL Fluor 610 or Quasar670 were ordered from LGC Biosearch Technologies. Probes were annealed prior to each experiment by hybridizing 6 µL primary probes, 3 µL secondary probes, 3 µL NEB3 buffer and 19 µL H_2_O in a thermocycler at 85°C for 3 min, 65°C for 3 min and 25°C for 5 min. The smiFISH protocol was adapted from Parker et al. (2020). In brief, animals were collected from plates containing many gravid adults and washed 3x in M9. Embryos were obtained by hypochlorite bleaching, washed 3x in M9 and transferred to 1.5 mL Eppendorf tubes. For fixation, embryos were resuspended in 1 mL acetone, submerged in liquid nitrogen for 1 min and incubated at -20°C for 10-60 min. Samples were treated with Stellaris Wash Buffer A (Biosearch Tech cat. SMF-WA1-60) containing 10% formamide (Invitrogen AM9342) for 5 min at room temperature (RT) and incubated with 100 µL Stellaris Hybridization Solution (Biosearch Tech. cat. SMF-HB1-10) containing 0.91 µL hybridized smiFISH probes at 37°C for 30 min with shaking at 450 rpm in the dark. Embryos were washed with Stellaris Wash Buffer A containing 2 µg/mL DAPI at 37°C for 30 min with shaking at 450 rpm in the dark. Finally, samples were incubated with Wash Buffer B at RT for 5 min, mounted in Vectashield mounting medium (Vector laboratories) and coverslipped.

### smiFISH combined with immunostaining

Primary and secondary smiFISH probes were hybridized as described for smiFISH. The experimental protocol was adapted from Tocchini et al. (2021) and carried out in quadruplicate. Gravid adults were splayed on a poly-L-lysine coated glass slide in 10 µL H_2_O. A coverslip was added and slides were transferred to liquid nitrogen. After a freeze crack, slides were immediately transferred to methanol at -20°C and incubated for 5 min. Slides were washed at RT in PBS (5 min), PBS with 0.5% Tween-20 (10 min and 20 min) and PBS (5 min). Samples were incubated in 50 µL fresh hybridization buffer (dextran sulfate (10% w/v) in one part formamide, one part 20×SSC and eight parts H_2_O) in a humidity chamber at 37°C for 1 hour. Hybridization buffer was removed and 50 µL hybridization buffer containing 1 µL pre-hybridized probes and primary antibodies (GFP goat polyclonal (Abcam #ab6673) 1/1000; RFP rabbit polyclonal (Rockland #600-401-379) 1/300) was added to the sample. Slides were incubated in a dark humidity chamber at 37°C for 6.5 hours and at 4°C overnight. Samples were washed twice with wash buffer (one part formamide, one part 20×SSC and eight parts H_2_O) and incubated with 100 µL wash buffer containing secondary antibodies (Alexa Fluor 488 donkey anti-goat (ThermoFisher Scientific #A-11055) 1/500; Alexa Fluor 555 donkey anti-rabbit (ThermoFisher Scientific #A-31572) 1/500) in a dark humidity chamber at 37°C for 1 hour. Slides were washed twice with wash buffer, mounted with Vectashield mounting medium (Vector laboratories) containing 0.7 µg/mL DAPI (Sigma-Aldrich D9542) and coverslipped.

### Immunostaining

The immunostaining protocol was based on Ramalho et al. (2020). Animals were cultured until plates contained many gravid adults and laid eggs. Larvae and adults were washed off using M9 and discarded. Approximately 1 mL of M9 was added to the plate to scrub off eggs with a gloved finger. Eggs were collected in 15 mL conical tubes and washed 3x with M9 and 1x with H_2_O. The majority of H_2_O from the wash step was removed and 10 µL of eggs suspension were transferred to a poly-L-lysine coated glass slide and freeze-cracked. Slides were immediately fixed at -20°C in P-buffer (3.7% formaldehyde, 75% methanol, 250 µL methanol, 250 µM EDTA and 50 mM NaF) for 15 min and methanol for 5 min. Embryos were dehydrated in a series of 90%, 60% and 30% ethanol at -20°C for 10 min each. Samples were washed 3x in PBS-Tween-20 before incubation with 50 µL blocking solution (1% bovine serum albumin and 10% serum in PBST) in a humidity chamber at RT for 1 hour. Blocking solution was wicked off and embryos were incubated with primary antibody (ERM-1 mouse monoclonal (DSHB Cat# ERM1, RRID:AB_10584795) 1/750) in blocking solution in a humidity chamber at 4°C overnight. Samples were washed 4x in PBST for 15 min and stained with secondary antibodies (Donkey anti-Mouse Alexa Fluor 488 (ThermoFisher Scientific #A-21202) 1/500) in blocking solution in a humidity chamber at RT for 2 hours. Unbound antibodies were removed by washing 3x in PBST for 10 min and 1x in PBS for 10 min. Finally, embryos were mounted using Prolong Gold Antifade with DAPI (Thermofisher) and coverslipped.

### Microscopy and image analysis

*C. elegans* larvae were paralyzed in a 10 mM Tetramisole solution in M9 buffer (0.22 M KH_2_PO_4_, 0.42 M Na_2_HPO_4_, 0.85 M NaCl and 0.001 M MgSO_4_) and mounted on a 5% agarose pad. For imaging of embryos, adults were splayed in M9 to release the embryos before mounting on a 5% agarose pad. Wide-field imaging was performed using an Axioplan2 upright microscope (Zeiss), equipped with a DIC polarizer and an Axiocam MRm CCD monochrome camera (Zeiss) and Plan-Apochromat 63x/1.4 Oil DIC M27 objective. Confocal imaging for smiFISH was performed using a Nikon Ti-U microscope equipped with Plan Apo VC 100x / 1.40 oil objective (Nikon, Japan) and Flash 4.0 v3 detector (Hamamatsu, Japan). Other confocal microscopy was performed using an Eclipse Ti2-E with perfect focus spinning disk (Nikon, Kōnan Japan) equipped with a CSU-X1-M1 confocal head (Yokogawa, Tokio Japan) and CFI Plan Apo λ 100x/1.45 oil objective. Images of the *erm-1*^*24xSunTag*^ strain were acquired using an Andor iXon DU-885 camera. A Photometrics Evolve EMCCD camera was used for heat-shock experiments, smiFISH combined with immunostaining and timelapse imaging of SunTag reporter strains. For all other experiments, a Prime BSI sCMOS camera was used. For quantifications, identical imaging approaches were applied within experiments. Microscopy data were acquired using Axiovision 4.x imaging software (wide-field microscopy) or Micromanager (confocal microscopy). Image processing was performed in FIJI in a non-destructive manner (Schindelin et al., 2012). Areas outside of the original field of view that became visible due to imaging rotation are indicated in white in the final figures.

### Quantification of localized translation

The distance of translation spots to the closest plasma membrane was calculated using Arivis Vision4D version 3.5.0. All analyzed embryos contained 40 to 60 cells and had a minimum of 25 translation spots. Embryonic cells were segmented based on the membrane marked by mCherry::PH and translation spots were detected by clustering of the scFv::GFP signal. The membrane signal was enhanced using a simple sharpening filter and Membrane Detection – Enhance Edges to ease segmentation of the individual cells. Detection of translation spots was improved using a discrete Gaussian denoising function to smooth out the signal and to filter out hot pixels. Embryonic cells were segmented using a Membrane-Based Segmenter and translation spots were segmented using a Blob Finder. The probability threshold of the blob finder operation was determined manually per embryo to achieve a correct segmentation of the translation spots. The split sensitivity of the blob finder operation was identical between *erm-1* and the randomized control but lower for the reporter to prevent over-segmentation. Everything outside of the embryos of interest and scFv::GFP signal within cells that were clearly dividing were discarded. Finally, the distance between the center of each translation spot and the closest edge of the membrane was measured in three dimensions (3D) in µm and exported via an Excel file. The number of translation spots within 0-0.35 micron from the membrane was divided by the total number of translation spots to calculate the proportion of localized translation. The randomized control was generated by flipping only the scFv::GFP signal with respect to the Z-axis to create a random distribution of translation spots while maintaining most translation spots within the embryo marked by the mCherry::PH. The exact same pipeline was followed in Arivis as the original data file to prevent computational bias.

### Quantification of overlap between translation spots and mRNA

To quantify the overlap between translation spots and mRNA, Z-stacks of embryos – stained for mRNA and protein (combined smiFISH and IF protocol) – were acquired at 0.4 μm intervals. Images were included for analysis if both mRNA and protein staining were homogeneous throughout the embryo. Maximum intensity projections of six Z-stack were generated using FIJI and background signal outside of embryos was discarded (Schindelin et al., 2012). FIJI ComDet v.0.5.5 was used to detect mRNA and translation spots in an unbiased manner and to calculate the integrated intensity of each spot (Katrukha, 2020). Only GFP spots with an integrated intensity >25000 were classified as translation spots and included in the analysis. The percentage of overlap was calculated by dividing the number of translation spots that overlap with mRNA by the total number of translation spots. Only embryos with >3 translation spots were used for analysis.

### Quantification of *erm-1* re-localization efficiency

An embryo with *erm-1*^*24xSunTag;20xPP7*^ + *pie-1p::pcp::mCherry::npp-9* background was imaged with 0.4 μm intervals over the Z-axis. Unbiased detection of translation spots was performed using the FIJI plugin ComDet v.0.5.5 (Katrukha, 2020). Duplicate Comdet results – due to detection of one translation spot over multiple Z-images – were discarded. The location of each translation spot was visually determined and categorized as “Nuclear pores”, “Plasma membrane” or “Cytosol”. Small differences in localization may be missed due to lower intensities of membranes orientated parallel to the Z-axis. For 4/154 translation spots the location could not be determined, which were discarded from analysis. The number of translation spots in each category was divided by the total number of translation spots to calculate the percentage.

### Quantification of membrane/cytoplasmic ERM-1 levels

Embryos of *erm-1::GFP* and *erm-1::1xGCN4*; *scFv::GFP* strains were imaged and analyzed using FIJI (Schindelin et al., 2012). Apical ERM-1 levels were obtained from the average of the peak intensity of three line scans perpendicular to the membrane between the ABa and ABp cells. Cytoplasmic ERM-1 levels were calculated from the average of the mean intensity of three cytoplasmic regions in the ABp cell. Apical ERM-1 levels were divided by cytoplasmic ERM-1 levels to calculate the ratio.

### Quantification of ACT-5 enrichment

Apical enrichment of YFP::ACT-5 was determined in FIJI by dividing the apical levels over cytoplasmic levels because of variable expression levels of the transgene. Analysis was performed in intestinal cells of comma to 1.5 fold embryos. Apical protein levels were calculated by averaging the peak intensity values of three line scans perpendicular to the apical membrane of intestinal cells of each animal. Cytoplasmic protein levels were obtained from the average of the mean intensity of three regions in the cytoplasm of intestinal cells per animal. Background levels were measured in three regions outside of embryos, averaged, and subtracted from the apical and cytoplasmic protein levels.

### Estimation of smiFISH probe binding efficiency

Probe binding efficiency for smiFISH was estimated using two different primary probes (against *24xGCN4* and *bfp*) targeting the same *24xGCN4::T2A::BFP::h2b* reporter mRNA, hybridized with different secondary probes (Quasar670 and CAL Fluor 610, respectively). The protocol for smiFISH and immunostaining was followed as described above. mRNA spots were detected using ComDet v.0.5.5 in FIJI (Katrukha, 2020; Schindelin et al., 2012). Data was exported to MS Excel to calculate the number of mRNA spots that overlapped between different channels. The number of overlapping spots was divided by the total number of mRNA spots for each channel to calculate the probe binding efficiency.

### Progeny size

The progeny size of WT, *erm-1*^*24xSunTag*^, *erm-1*^*24xSunTag;20xPP7*^ strains with or without re-localization by PCP::mCherry::NPP-9 was determined over the course of 72 hours from egg laying onset. Ten healthy looking animals were cultured at 20°C for every 24 hours before all laid eggs and larvae were counted and parental animals were transferred to fresh plates.

### Growth rate analysis

To collect synchronized L1 larvae, embryos were obtained by hypochlorite bleaching and hatched overnight in M9. Larvae were filtered through a 20 µm nylon net filter (Merck NY2004700) to discard unhatched eggs, and incubated at 20°C for 72 hours. At 0 hours (before incubation) and after 24 hours of incubation, animals were imaged using a Zeiss wide-field microscope. At 48 and 72 hours of incubation, animals were imaged using a S9i Digital Stereo Microscope (Leica, Wetzlar Germany) on 4x magnification. Images were analyzed in FIJI to calculate the body length as a measurement of growth (Schindelin et al., 2012). Animals were manually outlined and the plug-in Analyze Skeleton (2D/3D)(Arganda-Carreras et al., 2010) was used to determine the length of the worm. L1 larvae were omitted from the analysis if animals on the same plate had started egg laying.

### Morphology assessment

The intestinal morphology of embryos was studied in ACT-5::YFP containing strains. Animals were washed off plates using M9 and eggs were collected by scrubbing off plates using a gloved finger and M9. Intestines of 12 to 36 embryos, from stage E14 and older, were characterized and classified as “no defect” (no irregularities), “mild defects” (small structure irregularities or little widening of the lumen), “medium defects” (widened lumen, lumen extrusions, or small constrictions) or “severe defects” (significant disruptions). Similarly, larval morphological defects were examined by visual inspection in the ACT-5::YFP expressing strains. Intestinal morphology was characterized as “no defect” (no irregularities), “mild defects” (little widening of the lumen), “medium defect” (larger widening of the lumen or small constrictions) or “severe defects” (severe widening of the lumen or significant constrictions). Larval morphological phenotypes were characterized by E.v.d.S. and at least three blinded others, after which a consensus grade for each image was determined. If no consensus was reached (e.g. in case of a tie), the grade assigned by E.v.d.S was used (8/90 final grades).

### Statistical Analysis

Statistical analyses were performed using GraphPad Prism v.9. A D’Agostino-Pearson test was used to assess normality. For normally distributed data, comparisons of two populations were made using an Unpaired Student’s t-test (with Welch’s correction if the SDs of the populations differed significantly), and comparisons of >2 populations were done using a One-way ANOVA followed by a Bonferroni’s multiple comparison test. For data not drawn from a normal distribution, comparison of >2 populations was performed using a Kruskal-Wallis followed by a Dunn’s multiple comparison test. The used statistical tests and sample sizes are described in the figure legends. Sample sizes were not pre-determined by statistical tests. No data was excluded from analysis, unless described in the methods.

## Supplementary information

**Figure S1:**
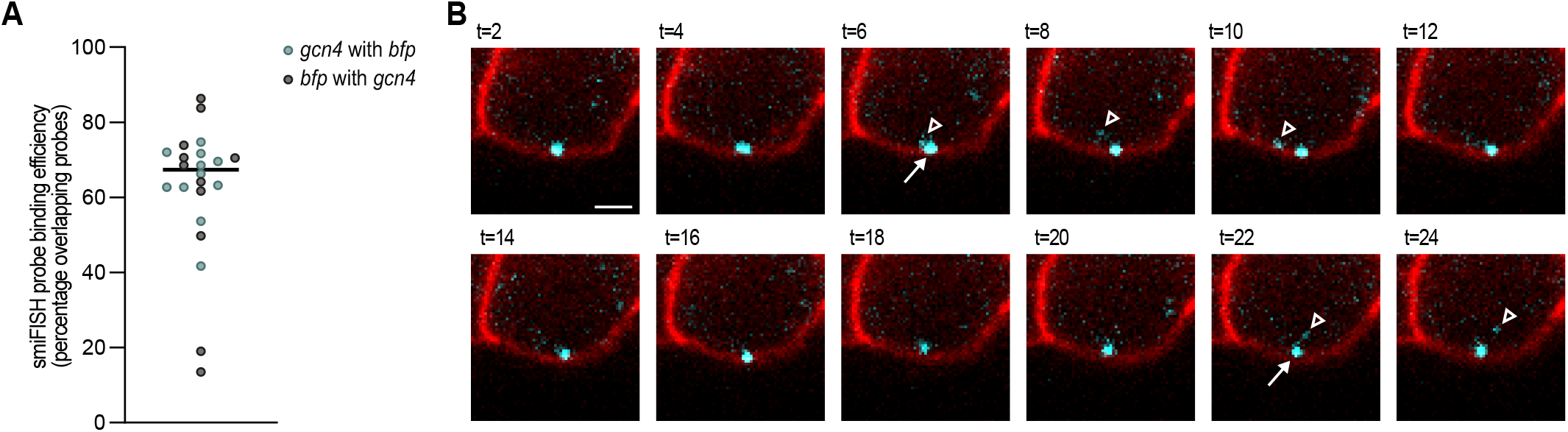
smiFISH efficiency and release of GCN4 proteins from translation spots. **(A)**smiFISH probe binding efficiency as measured by the overlap in *24xGCN4* and *BFP* smiFISH signals on *24xGCN4::t2a::BFP::h2b* mRNAs. Green dots represent the percentage *24xGCN4* smiFISH spots overlapping with *BFP* smiFISH spots, grey dots represent the percentage of *BFP* smiFISH spots overlapping with *24xGCN4* smiFISH spots, per embryo (n=11 embryos). **(B)** Time-lapse imaging of the release of mature protein from a translation spot over a 24 period. Depicted is a cell in an embryo expressing the SunTag antibody (scFv::GFP; depicted in cyan), a translation imaging reporter (*eft-3p::24xGCN4::AID::T2A::BFP::h2b::20xPP7::tbb-2 3’UTR*), the membrane marker (mCherry::PH; depicted in red) and the PCP protein (PCP::mCherry::PH; also depicted in red). The translation spot remained visible throughout the live imaging and is indicated by arrows. GCN4 proteins released from the translation site were transiently visible and are indicated by open trianglesGCN4. Scale bar: 1 µm.

**Figure S2:**
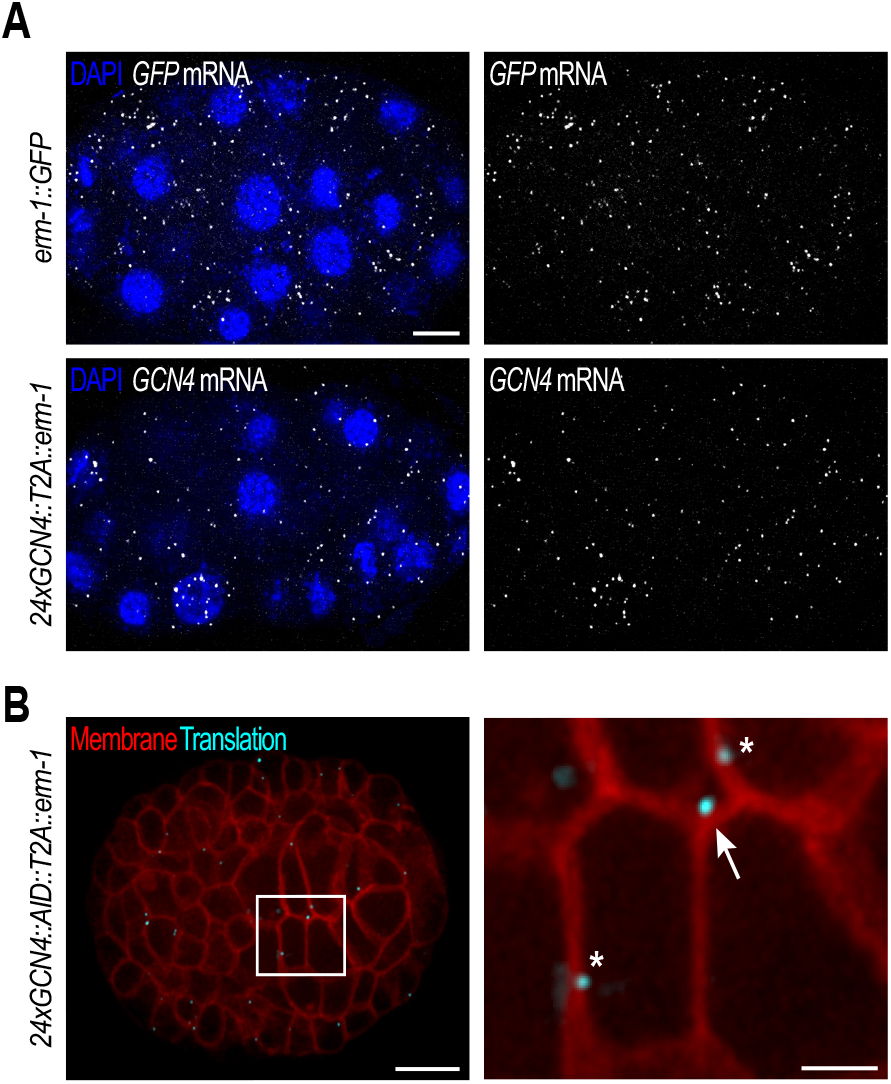
mRNA levels and localized translation in *erm-1*^*24xSunTag*^ embryos. **(A)**Representative images showing similar *erm-1* mRNA (smiFISH signal; gray) levels in embryos expressing endogenous ERM-1 tagged with GFP (*erm-1::GFP*; upper images) or embryos derived from the *erm-1*^*24xSunTag*^ strain. Nuclei are indicated by DAPI (blue). Scale bar: 10 µm. **(B)** Translation of *erm-1* (cyan) in close proximity of the basolateral (asterisks) and apical (arrow) membranes (red) of developing intestinal cells in a *24xGCN4::AID::T2A::erm-1* embryo. Scale bar: 10 µm (embryo); 2.5 µm (zoom-in).

**Figure S3:**
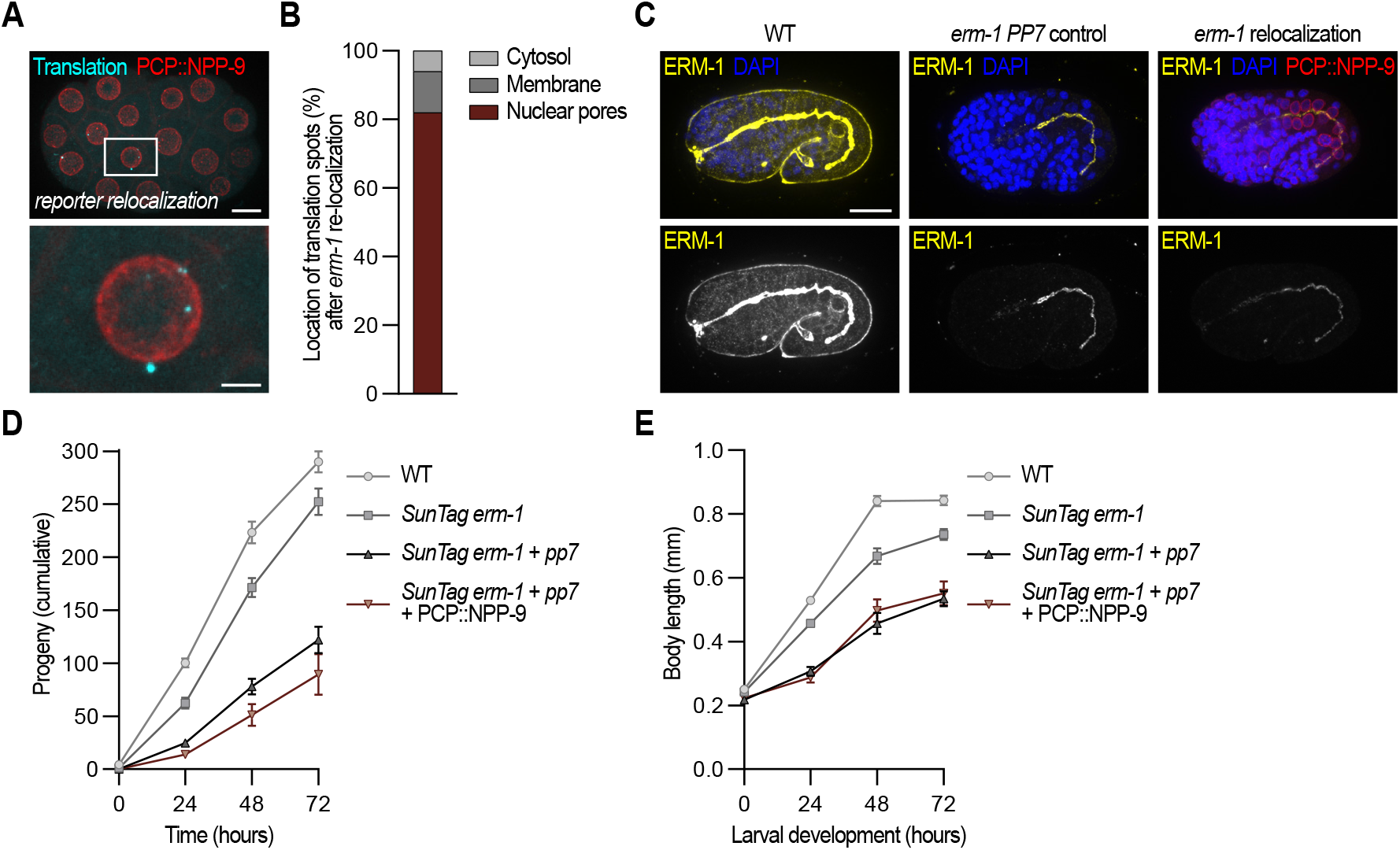
Re-localization of translation does not cause general phenotypes. **(A)** Re-localization of reporter mRNAs (*24xGCN4::AID::T2A::BFP::h2b::20xPP7::tbb-2 3’UTR*) to nuclear pore proteins (PCP::NPP-9; red) in the C. elegans embryo. Translation of reporter mRNAs is depicted by clustering of the SunTag antibody (scFv::GFP; cyan). Scale bar: 10 µm (embryo); 2.5 µm (zoom-in). **(B)** Quantification of the location of translation spots after *erm-1* re-localization to nuclear pores (n=150 translation spots). **(C)** Immunostained ERM-1 (yellow) in WT embryos (left images) or *erm-1*^*24xSunTag;20xPP7*^ embryos without (middle images) or with (right images) re-localization to nuclear pore proteins by PCP (PCP::mCherry::NPP-9; red). Nuclei are indicated by DAPI (blue). Scale bar: 10 µm. (D-E) Quantification of brood size **(D)** and larval growth **(E)** of WT strains, *erm-1*^*24xSunTag*^ strains and *erm-1*^*24xSunTag;20xPP7*^ strains with or without PCP::mCherry::NPP-9 expression. n=10 for each condition in (D), n=12 to 35 animals in (E).

**Table S3:**
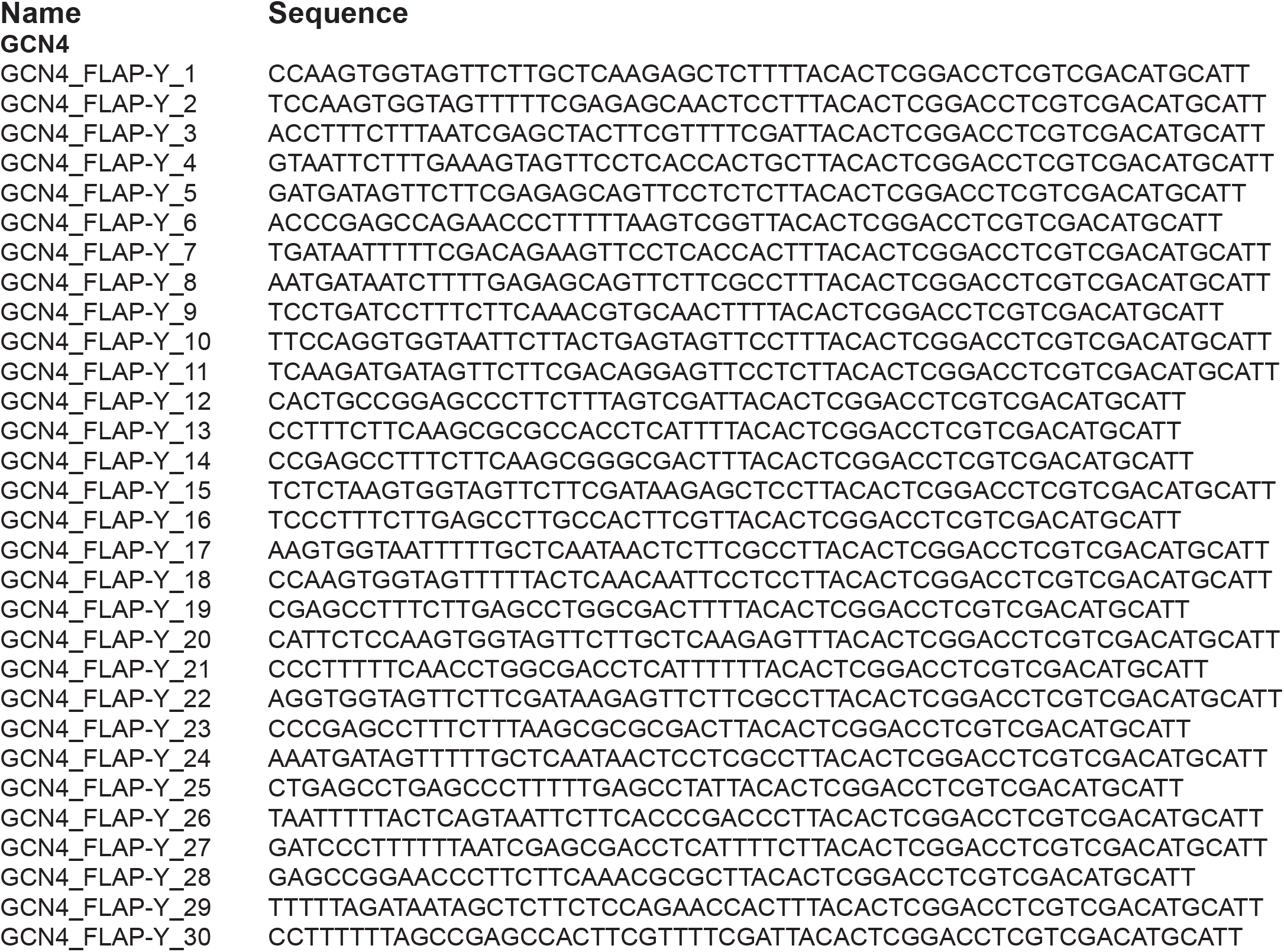
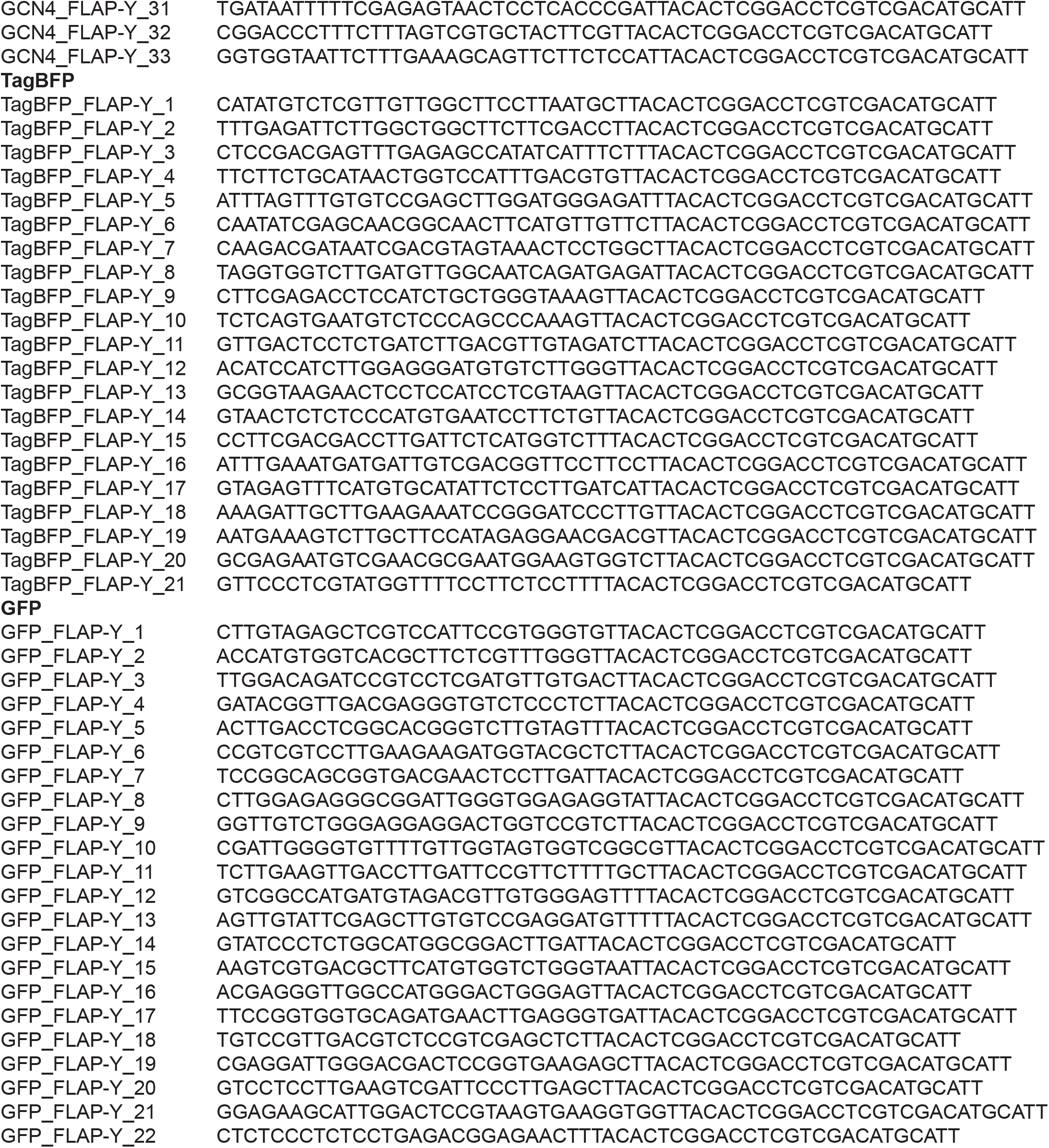
Primary probes used for smiFISH.

