## Supplemental Table 1 for "Localized translation of *erm-1* contributes to ERM-1 function in the *C. elegans* embryo"

| Strain | Genotype | Source | Method | Importan | Used in Figure |
|---|---|---|---|---|---|
| Box213 | erm-1(mib15[erm-1::eGFP]) I | Ramalho e | - | - | 1F, S2A |
| JM125 | caIs107[ges-1p::YFP::act-5] | CGC | - | - | 5I |
| PD1074 | C. elegans wildtype | CGC | - | - | 5F, S3C, S3D, S3E |
| SUR105 | ttTi5605(rubSi6[eft-3p::scfv(glo)::gfp(smu-1 introns)::tbb-2 3’UTR; 5' truncated gfp(smu-1 introns)::tbb-2 3’UTR]) II; par-3(rub33[1xgcn4::par-3]) III; tIs44[pie-1p::mCherry::PH(PLC1delta1) + unc-119(+)] | This paper | CRISPR/Ca | - | 1D |
| SUR106 | ttTi5605(rubSi6[eft-3p::scfv(glo)::gfp(smu-1 introns)::tbb-2 3’UTR; 5' truncated gfp(smu-1 introns)::tbb-2 3’UTR]) II; npp-9(rub34[1xgcn4::npp-9]) III | This paper | CRISPR/Ca | - | 1E |
| SUR119 | ttTi5605(rubSi6[eft-3p::scfv(glo)::gfp(smu-1 introns)::tbb-2 3’UTR; 5' truncated gfp(smu-1 introns)::tbb-2 3’UTR]) II; ltIs44[pie-1p::mCherry::PH(PLC1delta1) + unc-119(+)]; erm-1(rub39[24xGCN4::T2A::erm-1::20xPP7 in 3'UTR]) I | This paper | CRISPR/Ca | - | S3C |
| SUR120 | ttTi5605(rubSi6[eft-3p::scfv(glo)::gfp(smu-1 introns)::tbb-2 3’UTR; 5' truncated gfp(smu-1 introns)::tbb-2 3’UTR]) II; ltIs44[pie-1p::mCherry::PH(PLC1delta1) + unc-119(+)]; npp-9(rub37[PCP::mCherry::npp-9]) III; erm-1(rub39[24xGCN4::T2A::erm-1::20xPP7 in 3'UTR]) I | This paper | Cross SUR: | - | 5C, S3B, S3C |
| SUR125 | ttTi5605(rubSi6[eft-3p::scfv(glo)::gfp(smu-1 introns)::tbb-2 3’UTR; 5' truncated gfp(smu-1 introns)::tbb-2 3’UTR]) II; erm-1(rub39[24xGCN4::T2A::erm-1::20xPP7 in 3'UTR]) I | This paper | Cross SUR: | - | 5F, 5G, S3D, S3E |
| SUR126 | ttTi5605(rubSi6[eft-3p::scfv(glo)::gfp(smu-1 introns)::tbb-2 3’UTR; 5' truncated gfp(smu-1 introns)::tbb-2 3’UTR]) II; npp-9(rub37[PCP::mCherry::npp-9]) III; erm-1(rub39[24xGCN4::T2A::erm-1::20xPP7 in 3'UTR]) I | This paper | Cross SUR: | - | 5F, 5G, S3D, S3E |
| SUR127 | ttTi5605(rubSi6[eft-3p::scfv(glo)::gfp(smu-1 introns)::tbb-2 3’UTR; 5' truncated gfp(smu-1 introns)::tbb-2 3’UTR]) II; erm-1(rub42[erm-1::1xgcn4]) I | This paper | CRISPR/Ca | - | 1F |
| SUR130 | erm-1(rub6[24xGCN4::T2A::erm-1] I; caIs107[ges-1p::YFP::act-5] | This paper | Cross SUR: | - | 5D, 5E, 5H, 5I |
| SUR131 | erm-1(rub6[24xGCN4::T2A::erm-1] I; npp-9(rub37[PCP::mCherry::npp-9]) III; caIs107[ges-1p::YFP::act-5] | This paper | Cross SUR: | - | 5D, 5E, 5H, 5I |
| SUR132 | cxTi10816(rubSi12[eft-3p::24xGCN4::AID::t2a::tagbfp::h2b::tbb-2 3’UTR::20xpp7 *rubSi8]) IV; ttTi5605(rubSi6[eft-3p::scfv(glo)::gfp(smu-1 introns)::tbb-2 3’UTR; 5' truncated gfp(smu-1 introns)::tbb-2 3’UTR]) II; ltIs44[pie-1p::mCherry::PH(PLC1delta1) + unc-119(+)]; npp-9(rub37[PCP::mCherry::npp-9]) III; wrdSi23[eft-3p::TIF | This paper | CRISPR/Ca | not all ltIs4 | S3A |
| SUR154 | ttTi5605(rubSi6[eft-3p::scfv(glo)::gfp(smu-1 introns)::tbb-2 3’UTR; 5' truncated gfp(smu-1 introns)::tbb-2 3’UTR]) II; ltIs44[pie-1p::mCherry::PH(PLC1delta1) + unc-119(+)]; rubEx25[eft-3p::24xGCN4::t2a::tagbfp::h2b::tbb-2 3’UTR::72xpp7; myo-2p::tdtomato] | This paper | CRISPR/Ca | - | 2F |
| SUR54 | cxTi10816(rubSi8[eft-3p::24xGCN4::mIAA7 AID::t2a::tagbfp::h2b::tbb-2 3’UTR *rubSi7]) IV; ttTi5605(rubSi6[eft-3p::scfv(glo)::gfp(smu-1 introns)::tbb-2 3’UTR; 5' truncated gfp(smu-1 introns)::tbb-2 3’UTR]) II; ltIs44[pie-1p::mCherry::PH(PLC1delta1) + unc-119(+)]; wrdSi23[eft-3p::TIR1::F2A::mTagBFP2::AID*::NLS::tbb-2 3'UT | This paper | CRISPR/Ca | - | 1I, 1J, 1K |
| SUR57 | ttTi5605(rubSi6[eft-3p::scfv(glo)::gfp(smu-1 introns)::tbb-2 3’UTR; 5' truncated gfp(smu-1 introns)::tbb-2 3’UTR]) II; ltIs44[pie-1p::mCherry::PH(PLC1delta1) + unc-119(+)]; erm-1(rub6[24xGCN4::T2A::erm-1] I | This paper | CRISPR/Ca | - | 3B, 3C, 3D, 4A, 4B, 4C, S2A, S2B, S3D, S3E |
| SUR72 | ttTi5605(rubSi6[eft-3p::scfv(glo)::gfp(smu-1 introns)::tbb-2 3’UTR; 5' truncated gfp(smu-1 introns)::tbb-2 3’UTR]) II | This paper | CRISPR/Ca | - | 1C |
| SUR77 | ttTi5605(rubSi6[eft-3p::scfv(glo)::gfp(smu-1 introns)::tbb-2 3’UTR; 5' truncated gfp(smu-1 introns)::tbb-2 3’UTR]) II; ltIs44[pie-1p::mCherry::PH(PLC1delta1) + unc-119(+)]; erm-1(rub31[24xGCN4::AID::T2A::erm-1] I | This paper | CRISPR/Ca | - | S2B |
| SUR80 | cxTi10816(rubSi12[eft-3p::24xGCN4::AID::t2a::tagbfp::h2b::tbb-2 3’UTR::20xpp7 *rubSi8]) IV; ttTi5605(rubSi6[eft-3p::scfv(glo)::gfp(smu-1 introns)::tbb-2 3’UTR; 5' truncated gfp(smu-1 introns)::tbb-2 3’UTR]) II; ltIs44[pie-1p::mCherry::PH(PLC1delta1) + unc-119(+)]; rubSi13[pie-1p::PCP::mCherry::PH(PLC1delta1) *ltIs44] | This paper | CRISPR/Ca | not all ltIs4 | S1B |
| SUR81 | cxTi10816(rubSi12[eft-3p::24xGCN4::AID::t2a::tagbfp::h2b::tbb-2 3’UTR::20xpp7 *rubSi8]) IV; ttTi5605(rubSi6[eft-3p::scfv(glo)::gfp(smu-1 introns)::tbb-2 3’UTR; 5' truncated gfp(smu-1 introns)::tbb-2 3’UTR]) II; ltIs44[pie-1p::mCherry::PH(PLC1delta1) + unc-119(+)]; rubSi13[pie-1p::PCP::mCherry::PH(PLC1delta1) *ltIs44]; wr | This paper | CRISPR/Ca | not all ltIs4 | 2A, 2B, 2C, 2D, S1A |
