## Supplemental Table 2 for "Localized translation of *erm-1* contributes to ERM-1 function in the *C. elegans* embryo"

| Integration/mutation | guide RNA, type | guide RNA, sequence | (Repair) template, type | (Repair) template, sequence |
|---|---|---|---|---|
| *ttTi5605(rubSi6[eft-3p::scfv(glo)::gf* | sgRNA | ATGTCCTCCTGATTCCATGA | Plasmid (germline optimized version of pHR- | CACCTGACGCGCCCTGTAGCGGCGCATTAAGCGCGGC |
| *cxTi10816(rubSi7[eft-3p::24xGCN4::* | sgRNA | AGCTCAATCGTGTACTTGCG | Plasmid (24xGCN4 originally from pcDNA4T | CACCTGACGCGCCCTGTAGCGGCGCATTAAGCGCGGC |
| *cxTi10816(rubSi8[eft-3p::24xGCN4::* | crRNA | GCGGCCGCACCGGTGGTACC | PCR product | AAAAAGGGTTCTGGCTCGGGTCAGCGGCCGCACCGGT |
| *cxTi10816(rubSi12[eft-3p::24xGCN4* | crRNA | GCATTTATCGAATTCTTATT | PCR product (20xpp7 hairpins were a kind gi | GGGAACCAAGGCCGTCACCAAGTACACTTCATCCAAAT |
| *erm-1(rub6[24xGCN4::T2A::erm-1]* | crRNA | ACCCCAATCCATCTTCGACT | PCR product | CGTGAATTCTCACCGCTCACCCCAATCCATCTTCGACTC |
| *erm-1(rub31[24xGCN4::AID::T2A::e* | crRNA | GCGGCCGCACCGGTGGTACC | PCR product | AAAAAGGGTTCTGGCTCGGGTCAGCGGCCGCACCGGT |
| *erm-1(rub39[24xGCN4::T2A::erm-1:* | crRNA | GATATGACGAATACAAGAAG | PCR product | CGACGGTATCGATAAGCTTGATATCGCACTGACTACGA |
| *erm-1(rub42[erm-1::1xgcn4]) I* | 2xcrRNA | AAGACTCTCCGTCAAATCCG; ATATTG | PCR product | GCGAAACCGAAATATCGAAAAACCGAAAAACCTGCCG |
| *rubSi13[pie-1p::PCP::mCherry::PH(P* | crRNA | ATCAAATTTTCTTTTCCAGA | PCR product (PCP was a kind gift from the G | TCCCAAACAATTAAAAATCAAATTTTCTTTTCCAGATGA |
| *par-3(rub33[1xgcn4::par-3]) III* | crRNA | TTTCAGATCGATCATCATGT | ssODN | CATTTAATTTTTCTTTATATCTGCTCATTTTCAGATCGAT |
| *npp-9(rub34[1xgcn4::npp-9]) III* | crRNA | ATTCGAAAAGATTCACTTAC | ssODN | CCTGATATTTTTCATATTTCAGCTCTCCTTTCGCAATGGG |
| *npp-9(rub37[PCP::mCherry::npp-9])* | crRNA | ATTCGAAAAGATTCACTTAC | PCR product | CTGATATTTTTCATATTTCAGCTCTCCTTTCGCAATGGGT |
| *rubEx25[eft-3p::24xGCN4::t2a::tag* | - | - | Plasmid (72xpp7 hairpins were a kind gift fr | CACCTGACGCGCCCTGTAGCGGCGCATTAAGCGCGGC |
